# A unicellular walker controlled by a microtubule-based finite state machine

**DOI:** 10.1101/2021.02.26.433123

**Authors:** Ben T. Larson, Jack Garbus, Jordan B. Pollack, Wallace F. Marshall

## Abstract

Cells are complex biochemical systems whose behavior emerges from interactions among myriad molecular components. Computation is often invoked as a general framework for navigating this cellular complexity. However, it is unclear how cells might embody computational processes such that theories of computation, including finite state machine models, could be productively applied. Here, we demonstrate finite state machine-like processing embodied in cells using the walking behavior of *Euplotes eurystomus*, a ciliate that walks across surfaces using fourteen motile appendages (cirri). We found that cellular walking entails regulated transitions between a discrete set of gait states. The set of observed transitions decomposes into a small group of high-probability, temporally irreversible transitions and a large group of low-probability time-symmetric transitions, thus revealing stereotypy in sequential patterns of state transitions. Simulations and experiments suggest that the sequential logic of the gait is functionally important. Taken together, these findings implicate a finite state machine-like process. Cirri are connected by microtubule bundles (fibers), and we found that the dynamics of cirri involved in different state transitions are associated with the structure of the fiber system. Perturbative experiments revealed that the fibers mediate gait coordination, suggesting a mechanical basis of gait control.

## Introduction

Cells are complex physical systems controlled by networks of signaling molecules. Single cells can display remarkably sophisticated, animal-like behaviors ^1–3^, orchestrating active processes far from thermodynamic equilibrium in order to properly carry out biological functions ^4, 5^. Indeed, single cells can make decisions by sensing and responding to diverse cues and signals ^6^, execute coordinated movements ^7, 8^ and directed motility ^9–12^, and even solve mazes ^13, 14^ and possibly learn ^15–18^. Such behaviors in animals arise from neural activity and have been studied extensively, but we know comparatively little about the mechanisms of cellular behavior ^19, 20^. In individual cells, behaviors emerge directly through the joint action of chemical reactions ^21^, cellular architecture ^3^, physical mechanisms and constraints within the cell ^22, 23^, and interactions of the cell with its local environment ^24^. The links between information processing, decision making, and the physical manifestation of those processes as cell state transitions suggest that cellular behavior might be understood as an embodied computation ^25, 26^. The theory of computation has often been invoked as a general framework for understanding cellular dynamics ^25, 27–32^, environmental sensing by bacteria being a deeply studied example ^31–33^, and has been used to engineer programmable cell states ^34^. However, the manner and extent to which cells might embody functional, computational processes as well as the extent to which a computational perspective on cellular behavior might prove productive remains to be seen.

Among the microbial eukaryotes (protists), ciliates display some of the most striking examples of unicellular behavior, including hunting ^3^, sensorimotor navigation ^10^, and predator avoidance ^35^. Spirotrichous ciliates of the genus *Euplotes* are notable for their complex locomotion ^36–38^, using bundles of specialized cilia (cirri) to walk across surfaces ^36, 37^ (Figure 1A, Videos S1 and S2). Depending on the species, these cells generally have 14 to 15 ventral cirri arranged in a highly consistent pattern used for walking locomotion ^39^. *Euplotes* live in aquatic environments, and in addition to walking, use their cirri for swimming and rapid escape responses ^40^ (Video S2). Oral membranelles (Figure 1B) generate feeding currents to capture bacteria and small protistan prey and are also used for swimming. Early 20^th^ century protistologists were so impressed by the apparent coordination of cirri that they proposed the existence of a rudimentary nervous system, the neuromotor apparatus, to account for their observations ^38^. This theory was motivated by the presence of intracellular fibers connecting various cirri (Figure 1C), now known to be to be tubulin-based structures ^41, 42^.

**Figure 1.**
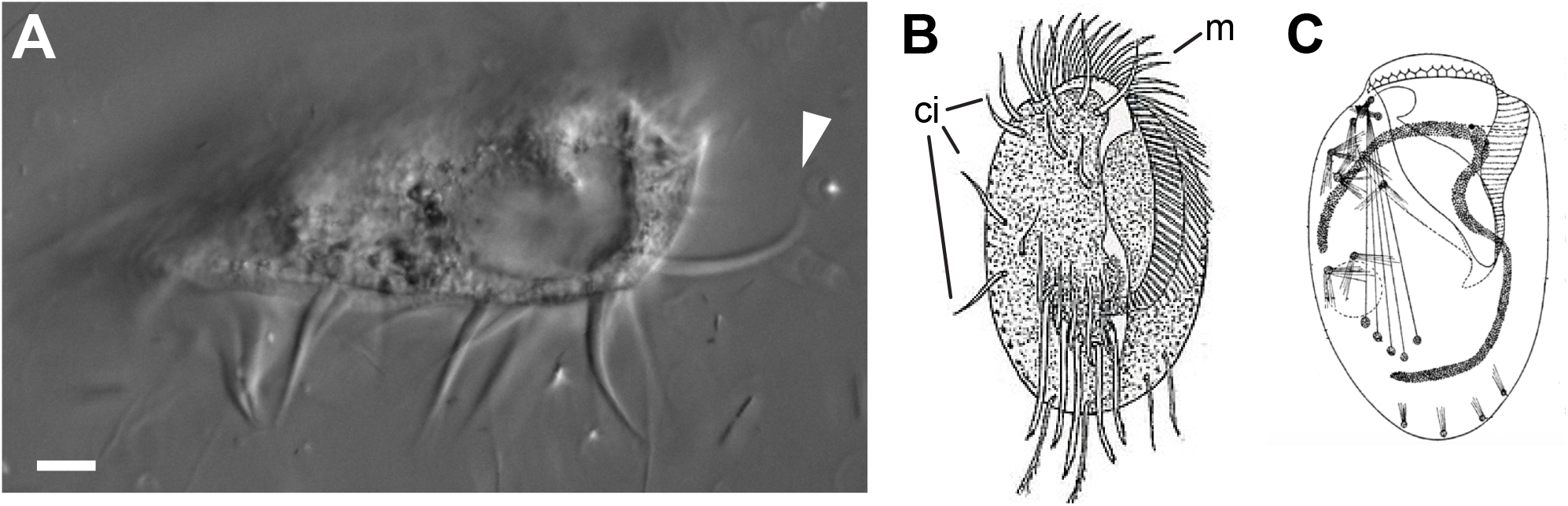
*Euplotes* exhibits highly polarized, complex cellular architecture and walks across surfaces using microtubule-based organelles called cirri, some of which are physically linked. (A) A single *Euplotes eurystomus* cell in profile displays its ventral cirri, which are used for walking locomotion across surfaces (arrowhead indicates a single cirrus stretching out from the cell). Scale bar is 10 µm. (B) A drawing of a *Euplotes* cell, viewed from the ventral surface, highlighting the complex, asymmetric structure of the cell. Notable features include the cirri (ci) and the membranellar band (m), wrapping from the top of the cell to the center, which is used to generate a feeding current to draw in prey items. Drawing adapted and obtained from Wikimedia Commons, from original source ^106^. (C) A drawing of a *Euplotes* cell, highlighting the fiber system associated with the cirri, historically referred to as the neuromotor apparatus. Drawing adapted from ^107^.

How can a single cell coordinate a gait without a nervous system? Although the walking movements of *Euplotes* are superficially similar to those of animals such as insects, the low Reynolds environment of aquatic microorganisms, where viscous forces dominate over inertial forces, imposes significant physical constraints on all movements that do not impinge on the movements of larger terrestrial animals ^43^. Coordination, to the extent that it exists in the gait of *Euplotes*, would require some kind of dynamical coupling among cirri or between cirri and some shared external influence. Recently, analytical techniques from statistical physics have been used to characterize, understand, and predict mesoscale dynamics in biological systems, including cellular behavior ^4, 5, 44, 45^. These approaches rely on coarse-graining the complexity of biological dynamics into states and analyzing the transitions among states. In particular, a state representation allows us to ask whether forward and reverse transitions among states are equal, a condition known as detailed balance ^4, 46^. Systems that violate detailed balance operate in a non-equilibrium mode, display net probability flows, and can produce directed cycles in state space^4, 47^. Broken detailed balance has been observed in the motility dynamics of cultured mammalian cells as well as the motility dynamics of a freely behaving flagellate protist ^5, 44^ and implies that non-equilibrium models are most applicable to such systems ^45^. Identification of broken detailed balance, therefore, highlights temporal irreversibility and can indicate active control of biological dynamics.

When information processing drives patterns of state transitions, such a system can be analyzed using automata theory, a fundamental level in the theory of computation ^48–50^. Automata theory can be used to address problems of decision-making and control in complex systems by providing predictive understanding that is independent of the underlying details of how a given process is implemented ^49^. Inspired by work considering cellular behavior in the context of the theory of computation ^25^, we hypothesized that walking cells might be governed by finite state automata with directed, processive movement arising from reproducible patterns of state transitions. Although the behavioral repertoire of *Euplotes* in different environmental conditions represents a rich and complex phenomenology involving information processing ^51^, for experimental tractability, we chose to focus on the reproducible, spontaneous linear walking behavior of cells. We reasoned that cellular walking might require some form of information processing to properly coordinate the movements of cirri.

The consistent structure of *Euplotes*, its mode of motility, and its ease of observation make these cells an ideal biological test-bed in which to apply theories of non-equilibrium statistical mechanics and embodied computation, both of which rely on describing a system in terms of discrete state transitions. Here, we use time-lapse microscopy and quantitative analyses to show that *Euplotes eurystomus* walks with a cyclic stochastic gait displaying broken detailed balance and exhibiting elements of stereotypy and variability, in accord with a finite state automaton representation. The observed dynamics are reminiscent of behavioral regulation in some cells and animals ^5, 52^ but contrast with many well-characterized examples of cellular and organismal motility ^7, 9, 10, 12, 53–58, 59^. Our results provide a demonstration of state machine-like processes governing cellular state transitions as well as an illustration of how such a computational perspective can drive mechanistic insight and serve as a framework for investigating the principles of behavioral control and non-equilibrium dynamics in single cells.

## Results

### A reduced state space is sufficient to describe walking dynamics

In order to ask whether cell behavior is governed by a finite state machine, we analyzed the walking behavior of *Euplotes eurystomus* cells, ^40^, focusing on the simplest case of uninterrupted, linear walking trajectories (Figure 2A,B, Video S1). Cells were placed onto coverslips on which free, spontaneous walking behavior was observed by microscopy (imaged at 33 frames/s). A focal plane at the cirrus-coverslip interface was chosen in order to clearly observe cirral dynamics (Figure 2A). The consistency of the relative spatial positioning of cirri across cells allowed us to give each of the 14 cirri an alphabetic label from a-n (Figure 2C). During walking, cirri move in a manner analogous to the recovery stroke-power stroke cycle executed by many eukaryotic flagella, first lifting off the substrate and sweeping close to the cell body before extending in roughly the direction of cell orientation before sweeping downward to reestablish contact with the substrate ^37, 60, 61^ (Figure 2B, Video S1). In each video frame, the walking state of the cell was encoded as a 14-element binary vector, with each element corresponding to a cirrus and receiving a value of “0” if the cirrus was in contact with the coverslip and stationary and a “1” if the cirrus was in motion or had moved in the preceding interval between frames (instances of stationary cirri held above the coverslip for a sustained period of time were not observed). The trajectories of 13 cells were manually tracked and annotated for a total of 2343 time points. This quantitative analysis revealed stepping-like cirral dynamics: cirri tend to undergo rapid movements followed by longer periods of quiescence (Figure 2D). Cirral dynamics seemed to lack any obvious patterns such as periodicity or repeating sequences of states (e.g. Figure 2D), implying that the state sequences are generated either by stochastic processes or complex deterministic mechanisms. This lack of periodicity (confirmed by autocorrelation analysis, Figure S1) or fixed phase relationships between appendage movements is different from those reported for various unicellular flagellates and the gaits of most animals ^58, 62, 63^.

**Figure 2.**
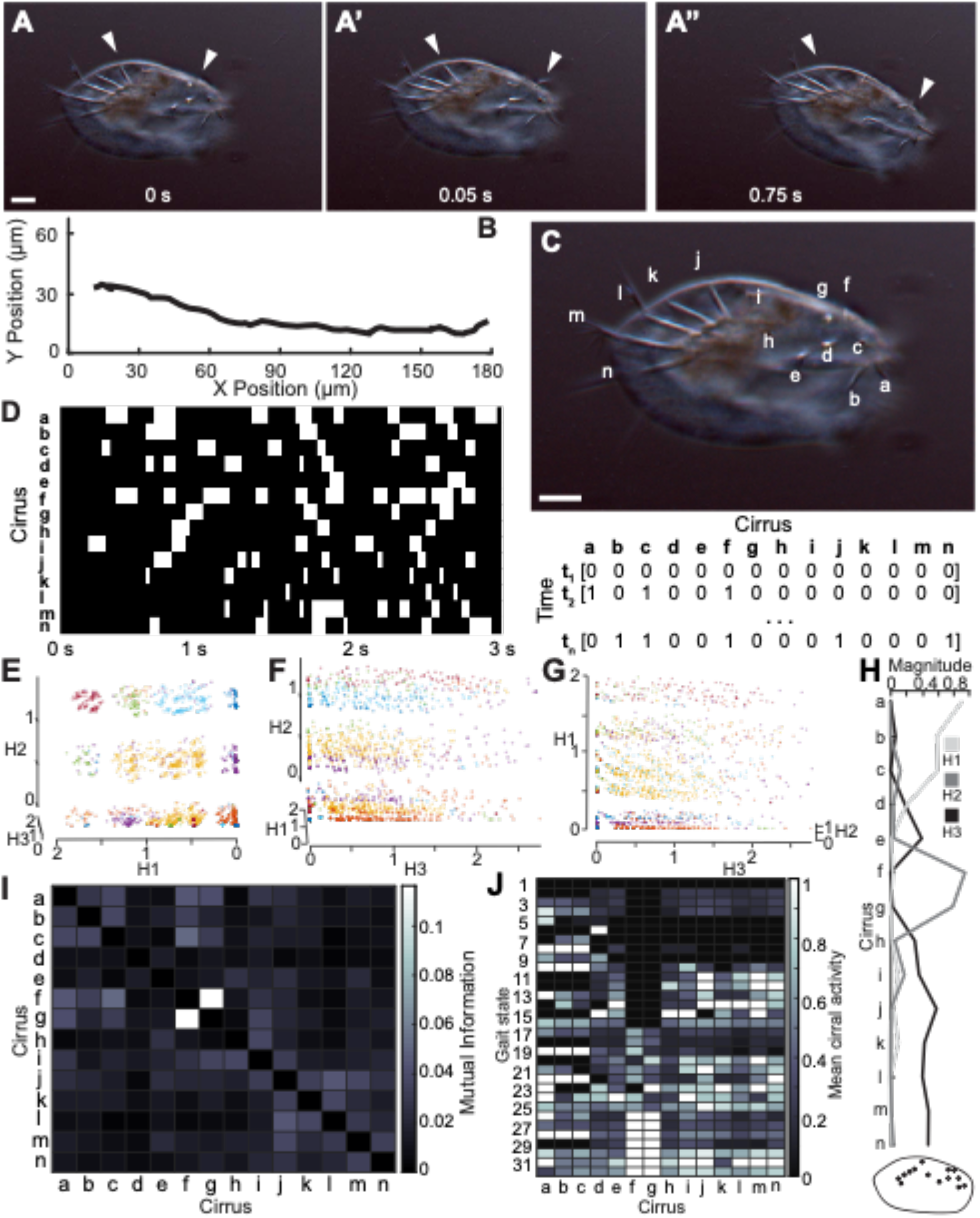
*Euplotes* walks across surfaces according to a complex gait, which can be described in a discrete, reduced state space with gait states corresponding to identifiable patterns of cirral activity. (A-A’’), The movements of cirri during walking locomotion are clearly visible by brightfield microscopy by focusing on a plane at the surface of the coverslip on which cells are walking. Three snapshots depict different time points during a single walking trajectory, and white arrowheads indicate cirri. In the panels from left to right, the cirrus indicated by the arrowhead on the left is stationary, stationary, and then moving, and the cirrus indicated by the arrowhead on the right is stationary, moving, and then stationary. Scale bar is 15 µm. (B) The trajectory of a cell during a single recorded trajectory as the cell walked across a coverslip from left to right. The cell position was manually tracked in each frame. (C) The scheme for encoding cirral dynamics during walking involved labeling each of the 14 distinguishable ventral cirri (a-n), and recording cirral activity in each frame, corresponding to timepoints (t_1_,…,t_n_), of recordings of walking cells as a 14-bit binary vector. Each entry in each vector is given a value of either 0 if the cirrus is not moving and in contact with the coverslip or 1 if the cirrus is moving. Scale bar is 15 µm. (D) Representative visualization of cirral dynamics for a single trajectory of a walking cell. These dynamics correspond to the walking trajectory in panel E. Each row corresponds to a cirrus and each column is a single video frame. White denotes cirral activity, a value of 1, in the vector encoding of dynamics from panel F. Note the dynamical complexity and discrete, stepping-like nature of cirral movements. (E-G) Three roughly orthogonal views of a plot displays the structure of all recorded cirral dynamics, encoded as in panel E, from 13 cells over 2343 timepoints in a reduced state space obtained by non-negative matrix factorization (NMF). Axes correspond to the components of the NMF (H1, H2, H3), and each point is a single timepoint. Randomized colors highlight the 32 clusters identified using the density-based spatial clustering of applications with noise (DBSCAN) algorithm ^70^. We refer to these clusters as gait states, and they correspond to unique configurations of cirral activity during walking locomotion. (H) A plot of the magnitudes associated with each cirrus corresponding to the components of the NMF of cirral dynamics shows distinct contributions from spatially distinct groups of cirri. Component H1, for example, is associated with activity in cirri a, b, and c. The tracing of a cell including the position of cirri has the same color map as the plot above and shows the grouping of the cirri corresponding to each component. (I) A heatmap of mutual information between all pairs of cirri shows that correlations in cirral activity correspond to the NMF components displayed in panel K. For example, cirri a, b, and c share mutual information with one another and are the cirri contributing to component H1. (J) A heatmap representation of the cirral activity associated with each of the 32 gait states. Values for each cirrus are the mean over all instances of the gait state. Note that each gait state has a unique signature of cirral activity. See also Figure S1, Video S1, and Video S2.

Despite the apparent complexity of cirral dynamics, we wondered whether there might be some underlying structure, which would allow us to effectively coarse-grain the dynamics in a principled manner. We first sought to obtain a reduced state space that could accurately describe the dynamics, as has proven successful in behavioral analysis of diverse living systems ^44, 45, 64–67^. Because our ultimate goal was to identify interpretable motifs among the patterns of cirral activity, which entail strictly nonnegative values, we chose to perform dimensionality reduction by non-negative matrix factorization (NMF), a technique that has been used to identify patterns for textual analysis, natural language processing, neural activity analysis, and gene expression analysis ^68, 69^. NMF revealed the cirral states to be well-described by a three-dimensional NMF space (Figure 2E-G, see Method Details and Figure S1 for more details). Factors of the NMF (H1, H2, H3) constitute the basis of the NMF space and correspond to non-overlapping groups of cirri. These factors indicate features of cirral activity that in various weighted combinations can be used to compactly represent all of the cirral activity measurements. These groups of cirri constitute spatially distinct partitions of cirri with respect to their positions on the cell body (Figure 2H). The dimensionality reduction of the gait state space arises in part from shared pairwise mutual information between groups of cirri (Figure 2I). Here, mutual information quantifies the amount of information about the patterns of activity of one cirrus gained by measuring the activity of another, and in this way acts like a generalized measurement of correlation. Therefore, dimensionality reduction by NMF reflects correlations in cirral activity.

Noting the apparent structure of the data visualized in the reduced gait state space in the form of clusters of points in NMF space (Figure 2E-G), we next sought to identify individual gait states defined in terms of similar patterns of cirral activity indicated by these clusters. To do this, we applied the density-based spatial clustering of applications with noise (DBSCAN) algorithm ^70^ to group the output of NMF into clusters in an unbiased fashion, with members of a given cluster sharing similar patterns of cirral activity (see Method Details and Figure S1 for more details). Although NMF can itself be used to define clusters, the advantage of clustering in two steps, NMF followed by DBSCAN, is that we could use the visual observation of clustering in the NMF output to confirm correct performance of the subsequent clustering step. The effect of this two-step approach to coarse graining, which is a commonly used in clustering of single-cell transcription data, is to reduce noise while retaining features that can accurately capture the complexity of cirral activity. Visual inspection in conjunction with silhouette coefficient (a metric of cluster cohesion and separation) analysis ^71^ revealed that 32 clusters accurately captured the visible structure in the reduced state space without overfitting (Figure 2E-G, Method Details and Figure S1). These reduced gait states correspond to distinct patterns of cirral activity (Figure 2J).

Taken together, our results reveal stereotypy in the spatiotemporal patterns of cirral activity, consistent with the behavior of a finite state machine. The discrete set of gait states, which exist in the reduced gait state space, demonstrate that cells make use of a subset of the possible patterns of appendage movement during walking locomotion.

### *Euplotes* walks with a cyclic stochastic gait

In order to relate the gait states identified in our cluster analysis to cell motility, we asked how changes in the number of active cirri may relate to cell movement. At low Reynolds number, velocity of a suspended particle is proportional to the difference between the net force acting on it and the opposing drag ^43^, so cell velocity provides a relative readout of net locomotive force. Naively, one might expect that the force associated with locomotion is roughly proportional to the number of moving appendages ^72^. Alternatively, the velocity might inversely correlate with the net change in cirral activity, which would be expected if stationary cirri were generating a pushing traction force as in crawling or climbing animals ^62, 73^ or if cirri execute a power stroke just before coming to rest as has been suggested previously ^37^. However, we found that data supported neither expectation: cell velocity was only weakly correlated with number of active cirri (R^2^=0.03), and instead, the largest cell velocities corresponded to small-to-moderate changes in the number of active cirri (Figure 3A). We hypothesized that transitions between gait states must be important to driving the forward progression of walking cells, and thus sought to determine whether such active coordination might manifest in the observed gait dynamics.

**Figure 3.**
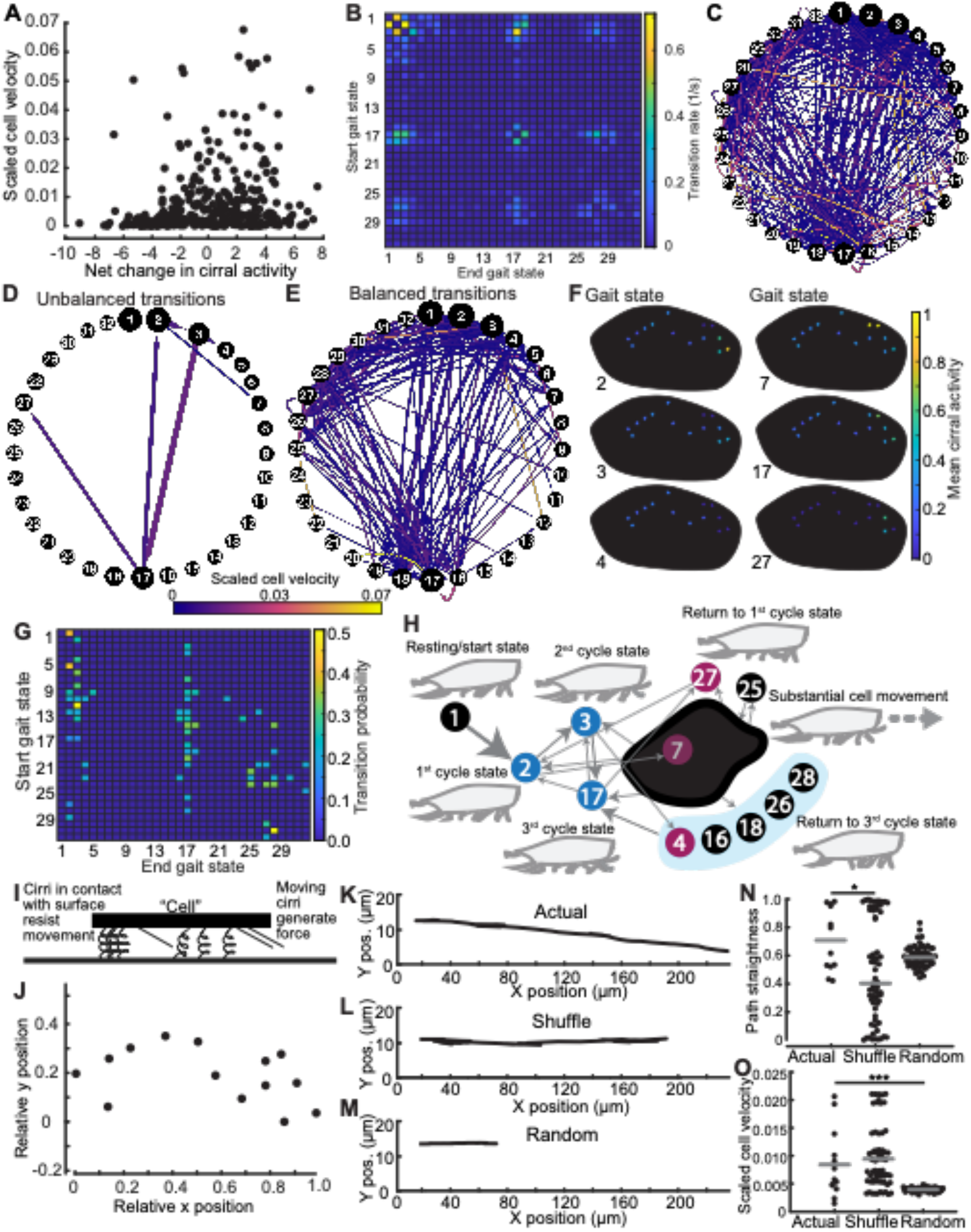
*Euplotes* walks with a cyclic stochastic gait exhibiting broken detailed balance, stereotypy, and state machine-like dynamics. All data for panels A-H are pooled from the walking trajectories of 13 different cells over 2343 timepoints and 1423 pairwise gait state transitions. (A) A plot of the mean net change in cirral activity versus the net scaled cell velocity associated with all transitions between the 32 gait states identified in Figure 2 shows that the change in number of active cirri is not strongly correlated with cell velocity (R^2^=0.03). Cell velocities were obtained from manually tracked walking trajectories and then scaled by dividing frame to frame displacements for each trajectory by the length of the cell being tracked and also dividing by the average frequency of cirral inactivity. Scaling provided a non-dimensional velocity scaled by natural units of the system. Note that at low Reynolds number, velocity should be proportional to force ^43^, so this plot also reflects the net walking force generated by the cell. Net change in cirral activity was computed using the data presented in Figure 2. Note that the largest velocities are associated with small negative and small to moderate positive net changes in cirral activity. (B) The transition matrix of all gait state transitions, with rows representing the starting state and columns indicating the ending state, exhibits broken detailed balance. Rates were estimated by dividing the total number of observed transitions between each state pair and dividing by the total time observed. Under detailed balance or equilibrium conditions, transitions from one state to another should be balanced by reverse transitions. Lack of this kind of reversibility, as seen by the lack of symmetry of the heatmap across the diagonal, indicates broken detailed balance and non-equilibrium dynamics. (C) A directed graph representation of all gait state transitions. Nodes correspond to the 32 gait states, and node sizes are scaled by the proportion of total time cells spent in each state. Directed edges are represented by arrows between nodes and signify state transitions. The size of the arrows is scaled by transition rates as in panel b. Edge color represents scaled cell velocity as in panel A according to the indicated color scale. (D) A subset of transitions visualized as in panel C shows the restricted and relatively high frequency nature of unbalanced, non-equilibrium-like transitions. Only transitions that were observed to happen more than one time and exhibiting a significant difference between forward and reverse transitions (p<0.05 by binomial test, see Method Details) are displayed. (E) A subset of transitions, similarly to panel d, except that only the balanced transitions, lacking a significant difference between forward and reverse transitions (p<0.05 by binomial test) are displayed, also show a complex and widespread structure, this time of balanced, equilibrium transitions. Note that the majority of transitions associated with high cell velocity involve equilibrium-like dynamics. (F) Examples illustrating the spatial organization of cirral activity corresponding to gait states. Some states, such as 7, correspond to activity in spatially discrete groups of cirri, while others, such as 17, correspond to cirral activity across the cell. The gait states displayed here are those involved in unbalanced transitions. (G) A heatmap of transition probabilities between states, showing only the most probable transitions from a given state with all others set to zero, shows distinct structure. In cases were multiple state transitions from a state were tied for the highest probability, all of these transitions are displayed. Fewer than half of the total states are recipients of multiple high probability transitions, and many states are the recipients of no high probability transitions. (H) A representation of functional states and transitions between them highlights the machine-like nature of the gait of *Euplotes*. Gait states are represented as circles with numerical labels. Blue circles represent states that are both recipients and sources of unbalanced transitions as identified in panel D and constitute the three cycle states. Red circles represent states that are recipients but not sources of unbalanced transitions as identified in panel D. Black circles correspond to gait states that are associated only with balanced transitions as in panel E. States receiving no more than one unique high probability transition from states with only a single highest one as identified in panel g were grouped together into a compound state represented by the dark gray blob. The blue background behind states 4, 16, 18, 26, and 28 indicates that these states all share the same highest probability transitions between states identified in this panel, and thus, the group constitutes a single compound functional state. Arrows represent the highest probability transitions between the states, including compound states composed of multiple gait states as identified in Figure 2 (dark gray blob and blue background). Gait state 1 is also depicted, as it is the state in which cells spent the most time over all walking trajectories and also is uniquely the state from which cells begin walking. Cells also frequently return to the state during walking. Further, transitions from gait state 2 from gait state 1 constitute the single highest frequency transition. Together, all identified states in this panel constitute functional states. Arrows represent the most probable transitions between functional states, and all unbalanced transitions are also represented with size scaled by their proportional probability compared to all other transitions emanating from the source functional state. Cartoons are a walking cell in profile with cirri in a configuration representative of the corresponding functional state. Labels refer to the apparent functional role of states and their associated transitions. Beginning from gait state 1, the resting/start state, cells are most likely to follow transitions from gait state 2 to 3 to 17 at which point cells are likely to enter the functional state associated with substantial cell movement involving variable balanced transitions between a number of gait states. Transitions are then likely to lead back toward the cycle states. Note that while this representation of gait dynamics highlights the most probable transitions, substantial variability, primarily involving reversible transitions, occurs during walking trajectories. (I) A cartoon depicts the basic features of our model of a walking cell. Cirri generate a constant force when moving and resist displacement in a spring-like fashion when in contact with the substrate. Force and torque balance yield cell displacement and orientation in each simulation timestep. (J) A plot of the relative average resting cirral-surface contact positions measured from the 13 cells used for gait analysis, which were used for all simulations and define simulated cell position and orientation. (K) A representative trajectory of a simulation of a walking cell using experimentally recorded cirral activity illustrates the linear, processive motion, which matches experimentally measured cell walking dynamics qualitatively and semi-quantitatively. (L) A representative trajectory of a simulation of a walking cell using the same experimentally recorded cirral activity as in panel K but in a shuffled order illustrates a decrease in processivity of cell movement. (M) A representative trajectory of a simulation of a walking cell with random patterns of cirral activity with the same average level of cirral activity and the same number of timesteps as in panels k and l illustrates the decrease in cell speed. (N) A plot of path straightness measured from simulations using actual cirral pattern measured experimentally (Actual), actual cirral patterns in a randomly shuffled order (Shuffle), and randomly generated cirral activity with the same average cirral activity as actual patterns (Random) suggest that the order of gait state transitions matters for processivity of cell movement due to a predicted decrease in path straightness for Shuffle trajectories. Path straightness is the distance between the start and end of the cell trajectory divided by the sum total path length. The trajectories of the 13 cells analyzed for Figure 2 were used for the Actual simulations. For the Shuffle simulations, cirral activity patterns from each of the 13 cells were shuffled to generate a random order five times. For the Random simulations, five sets of patterns of lengths equal to each of the 13 cell trajectories were generated. The asterisk indicates p=0.04 by the Wilcoxon rank sum test. (O) A plot of scaled cell velocity for the simulations described for panel k predict that the gait states are responsible for generating cell velocities as indicated by decreased scaled cell velocity for random patterns of cirral activity. Scaled velocity was computed as described for panel a and averaged over the entire cell trajectory. Three asterisks indicates p<0.001 by the Wilcoxon rank sum test. See also Figure S1 and Figure S2.

To search for evidence of active coordination dictating gait state transitions, we calculated the forward and reverse transition rates between states from the 1423 pairwise transitions in our dataset as *N_ij_*/*T* where *N_ij_* is the total number of transitions observed between states *i* and *j*, and *T* is the total observation time (Figure 3B, see Method Details). We note that the transition rate we define here is proportional to the probability current *J_ij_* = P_i_k_ij_ where P_i_ is the probability of state i and k_ij_ is the conditional transition probability analogous to a chemical reaction rate. Although walking, and cilia-based motility in general, is fundamentally non-equilibrium in the sense that it requires energy input, a coarse-grained representation of motility such as the one we pursue here need not exhibit broken detailed balance ^5^. However, broken detailed balance in gait state transitions (unbalanced forward and reverse transitions) would indicate a non-equilibrium process dictating temporal directedness (irreversibility) to gait state transitions, thus implicating active coordination in terms of stereotyped sequential changes in cirral activity. Importantly, broken detailed balance in gait state transitions define an identifiable temporal direction in sequences of gait state transitions. The presence of strongly unbalanced transitions such as from gait state 3 to 17 versus 17 to 3 suggested broken detailed balance, and indeed, a number of forward and reverse transitions were found to be significantly unbalanced by the binomial test (see Method Details). Unbalanced transitions can also arise in equilibrium systems that have not yet reached steady state, in which case transition rates may change in time. To test whether the gait operates at steady state, we checked whether the total number of transitions into each state were balanced by total transitions out of that state: **∑_j_ *N*_ij_ = ∑_j_ *N*_ji_**. Consistent with steady state dynamics, we found that this condition held to within a difference of at most a single transition. Although there are cases where non-equilibrium state transitions can nevertheless exhibit balanced transitions, the presence of unbalanced transitions existing as part of loops in state space unambiguously indicated non-equilibrium dynamics. To illustrate this non-equilibrium or temporally irreversible character of gait state transitions, we apply the Kolmogorov criterion, which specifies the necessary and sufficient condition for reversibility that the product of transition probabilities traversing any closed loop in state space must equal the product of the transition probabilities in the reverse direction ^5, 74^. Due to the presence of unbalanced transitions, the gait of *Euplotes* clearly violates this condition; for example, *k*_1,2_*k*_2,3_*k*_3,17_*k*_17,1_ = 0.003 ≠ *k*_1,17_*k*_17,3_*k*_3,2_*k*_2,1_ = 0.0005, where each *k_i,j_* is the conditional probability of transitioning from state *i* to state *j* estimated as *N_ij_*/*N_i_* with *N_i_* the total number of transitions from state *i*. To further understand the degree to which detailed balance was broken, or, similarly, the distance from equilibrium, we calculated the entropy production rate ^5^. Following the procedure detailed by Wan and Goldstein ^5^, we obtained a lower bound estimate for an entropy production rate of 0.4 nats, similar to the value reported for strongly non-equilibrium gait transitions observed in a flagellate ^5^. Walking *Euplotes* cells, therefore, have a non-equilibrium gait, displaying temporal order in sequences of appendage movement, despite a lack of standard gait periodicity.

We next sought to better understand the organization and sequential logic of this unusual gait. First, we noted that gait state transitions appear constrained: only 322 of the 1024 possible types of transitions were observed to occur at least once, and within this restricted set, only 173 occurred more than once (Figure 3C). Crucially, the presence of broken detailed balance revealed directed cycles of cirral activity during locomotion pushing the system away from equilibrium, yielding a temporal directedness or irreversibility to sequences of gait state transitions in spite of the lack of standard gait periodicity. To investigate the nature of these cycles, we grouped transitions into two categories: balanced transitions, which satisfy detailed balance, and unbalanced transitions, which do not (see Method Details). This partitioning allowed us to separately investigate unbalanced, non-equilibrium-like and balanced, equilibrium-like transitions (Method Details, Figure 3D,E). We found that unbalanced transitions occur at relatively high frequency but involve a small number of states (Figure 3D). Of the 32 gait states, only states 2, 3, 4, 7, 17, and 27 (Figure 3F) are associated with unbalanced transitions, and among these states, three formed a directed cycle following 2➔3➔17➔2. We had expected that unbalanced transitions might be associated with a “power stroke” (in the sense of occurring simultaneously with cell movement) but found instead that high cellular velocities tend to be associated with balanced transitions (Figure 3D,E) and that relatively few transitions corresponded to substantial cell movement (Figure 3C). Additionally, we found that, with the exception of transitions between states 1 and 2, the highest frequency transitions are unbalanced (Figure 3D).

Notably, the most frequent balanced transitions were associated with transitions into and out of gait state 1, a unique “rest state” which involves no cirral movement (Figure 2J). Furthermore, we found by autocorrelation analysis of gait trajectories that gait state 1 uniquely exhibited significant positive autocorrelation (see Method Details, Figure S1). Investigation of transitions to and from gait state 1 revealed that although transitions between gait states 1 and 2 are balanced, the most frequent transitions out of gait state 2 are strongly biased toward transitions into gait state 3, from which other strongly biased transitions also frequently occur, including the cycle of biased transitions mentioned above. Despite the presence of high frequency unbalanced transitions, the gait of *Euplotes* involves highly variable trajectories through gait state space. The picture of walking trajectories that emerges is one of stochastic excursions from gait state 1 into non-determinate paths through state space involving a mix of balanced and unbalanced transitions. The majority of cell movement occurs during infrequent, equilibrium-like (balanced) transitions. These balanced transitions are temporally reversible, but we wondered whether there might be some asymmetry in the amount of cell movement due to transitions leading toward versus away from the cycle states. We found that although balanced transitions leading away from the cycle states account for 7.6% more cell movement, the difference in these transitions is not statistically significant (p=0.09 by two-sample Kolmogorov-Smirnov test). Temporal irreversibility, or directedness, in sequential changes in cirral activity arises from biased, non-equilibrium-like (unbalanced) transitions, occurring at relatively high frequency from a small subset of states. A natural question arising from the observation of these complex patterns is whether gait state transitions conform to a first-order Markov process, often referred to as “memoryless” ^75^, which entails that transition probabilities are determined completely by the present state, and that previous dynamics contribute no additional predictive information ^76, 77^. Does the cell retain some “memory” of past cirral activity that might influence future cirral activity? While not conclusive, several analyses (see Method Details and Figure S1) suggested that *Euplotes* retains some “memory” of the prior sequence of cirral movements during locomotion in that the gait may not conform to a continuous- or discrete-time first-order Markov process.

Taken together, our analysis revealed a mixture of unbalanced transitions driving cycles and balanced transitions arranged as networks, for which we propose to apply the term “cyclic stochastic gait”. The cyclic stochastic gait of *Euplotes eurystomus* incorporates elements of both stereotypy and variability in gait dynamics, in terms of biased transitions and non-determinate sequences of gait state transitions respectively. Forward progress of the cell is not produced merely by a physical ratchetting process driven by unpatterned fluctuations in cirral activity, nor is it produced by a highly regular, deterministic process like a clock. It has been argued that significant computation arises in physical systems exhibiting such a mix of stereotypy and variability ^48, 78–80^ in the sense that the time-evolution of the system is most compactly described as the result of a computational process involving state transitions, memory, and decision rules, rather than periodic oscillations or random coin flips.

Our analysis suggested a computational underpinning of gait, so we sought to better understand the functional organization of the dynamical patterns, the sequential logic of the gait. Focusing on the dominant structure of gait transitions in terms of transition probabilities (Figure 3G, Method Details) allowed us to derive a simplified, functional representation of stereotypy in gait dynamics as depicted in Figure 3H. We found that few states were the recipients of the majority of the highest probability transitions and that many received no high probability transitions (Figure 3G). Additionally, we found a “cloud” of states linked by low-probability balanced, equilibrium-like fluctuations. Nearly all of the states receiving high probability transitions were either the three “cycle states” or else fed cycle states with their highest probability transitions, with the majority feeding gait state 17. Although gait state 1 is not the recipient of any individual high probability transitions, we identified it as the unique “start” state from which cells initiate walking. Beginning with this start state, cells transition with high probability to gait state 2, also one of the highest frequency transitions and the first state in the 2➔3➔17➔2 cycle of unbalanced transitions. From this first cycle state, cells transition to gait state 3, the second cycle state, with highest probability and frequency and then similarly on to gait state 17, the third cycle state. This sequence from the start state through the cycle states corresponds to increasing amounts of cirral activity. Although the highest probability transitions from the third cycle state to any single gait state tend to return to the first or second cycle state with equal probability, cells in fact transition to the equilibrium “cloud” of motility-associated states with overall higher probability. Return to the cycle states tend to occur through various moderately high probability transitions from the motility state cloud or through a restricted set of intermediate states. In conjunction with this set of transitions, we also noted unbalanced transitions stemming from the cycle states to the motility state as well as the presence of intermediate states from a given cycle state that subsequently feed the next cycle state.

Altogether, the picture of stereotypical gait dynamics that emerges is of biased transitions involving cycle states preceding relatively low probability, unbiased transitions associated with substantial cell movement before returning to the start or cycle states and beginning the sequence again. While this general sequence is repeated during walking, there is considerable variability or apparent stochasticity in the details of gait state transitions departing from the start state with increasingly variable transitions as any given sequence progresses. Given that the motion of the cell alternates between periods of highly processive forward motion and other periods in which the cirri seem to flail ineffectually without driving the cell forward, one might assume that the periods of effective motion would correspond to the highly directed cycle of unbalanced transitions and the periods of flailing would correspond to the cloud of balanced, random transitions, but in fact the opposite is the case. We hypothesized that sequences involving the cycle states serve to establish configurations of cirri necessary for cells to later transition between states from which substantial forward progress of the cell is generated. Many state transitions along any instance of the stereotyped sequence are unbiased; however, biased, high probability transitions, presumably resulting from active cellular control, give temporal irreversibility to the sequence. We note here that although our coarse-graining procedure to identify gait states does not constitute a unique representation of gait structure, we can be confident that our analysis captures structure in gait dynamics, the presence of broken detailed balance in particular. In general, coarse-graining of the state space of a system can obscure broken detailed balance, but the net flux of transitions in the state space of a system should not arise in an illusory fashion based on a coarse-graining procedure or partial observation of a system ^81–83^. Additionally, we have chosen a coarse-graining procedure based on the properties of our particular data and demonstrate its performance on simulated cirral dynamics with varying noise and for the case of unpatterned, random cirral fluctuations (Figure S1, Method Details).

Finally, we sought to investigate the functional significance of the sequential logic of the gait in driving processive cell movement. To do so, we developed a simple model based on a coarse-grained physical picture of cellular walking (Figure 3I). Briefly, we consider a 2D system where a cell walks using its 14 cirri, which can exist in two states: moving or not moving. As we are agnostic to the details by which cirri generate forces involved in motility, we model cirri as producing constant force in the direction of cell orientation while moving and resisting displacement by acting as linear springs when not moving. Cell position and orientation is defined in terms of the equilibrium positions of the cirri. For simulations, we use relative positions of cirri taken from stationary cell measurements (Figure 3J). In each timestep of the simulation, we calculate a cell displacement based on the sum of forces due to the cirri and changes in cell orientation from the sum of torques (see Method Details). We found that this simple model was sufficient to qualitatively and semi-quantitatively reproduce the linear trajectories of walking cells when we ran simulations using the actual patterns of cirral activity from walking cells (Figure 3K, Figure S2). When we ran simulations with either the same gait states as those from actual cells but in a shuffled order or random cirral activity with the same average cirral activity as actual cells, we found that path straightness significantly decreased in the case of shuffled transitions (p=0.04 by Wilcoxon rank sum test) and scaled cell velocity significantly decreased in the case of random activity (p=0.003 by Wilcoxon rank sum, Figure 3K-O). Note that in both the case of shuffled transitions and random cirral activity, gait state transitions satisfy detailed balance (Figure S2). These results suggest that the sequential logic of the gait is functionally significant in a manner consistent with our biophysical picture of walking.

### The microtubule-based fiber system mediates gait coordination

The complex yet sequentially structured gait patterns in conjunction with our simulation results are consistent with the existence of some form of nontrivial gait coordination. What physical machinery could embody the information processing required to generate the stochastic cyclic state transitions seen during *Euplotes*’ walking? We reasoned that there must be some form of coupling or communication between cirri or feedback between gait states and cirral dynamics. Since the early 1900s, the role of the system of cytoskeletal fibers associated with cirri as conduits of information between cirri during cellular locomotion, supported by microsurgical experiments, has been a dominant yet contentious hypothesized mechanism of gait coordination ^84–86^. We wondered whether the structure of the cytoskeletal fiber system associated with cirri (Figure 4A) could give some insight into how cirri might be coordinated.

**Figure 4.**
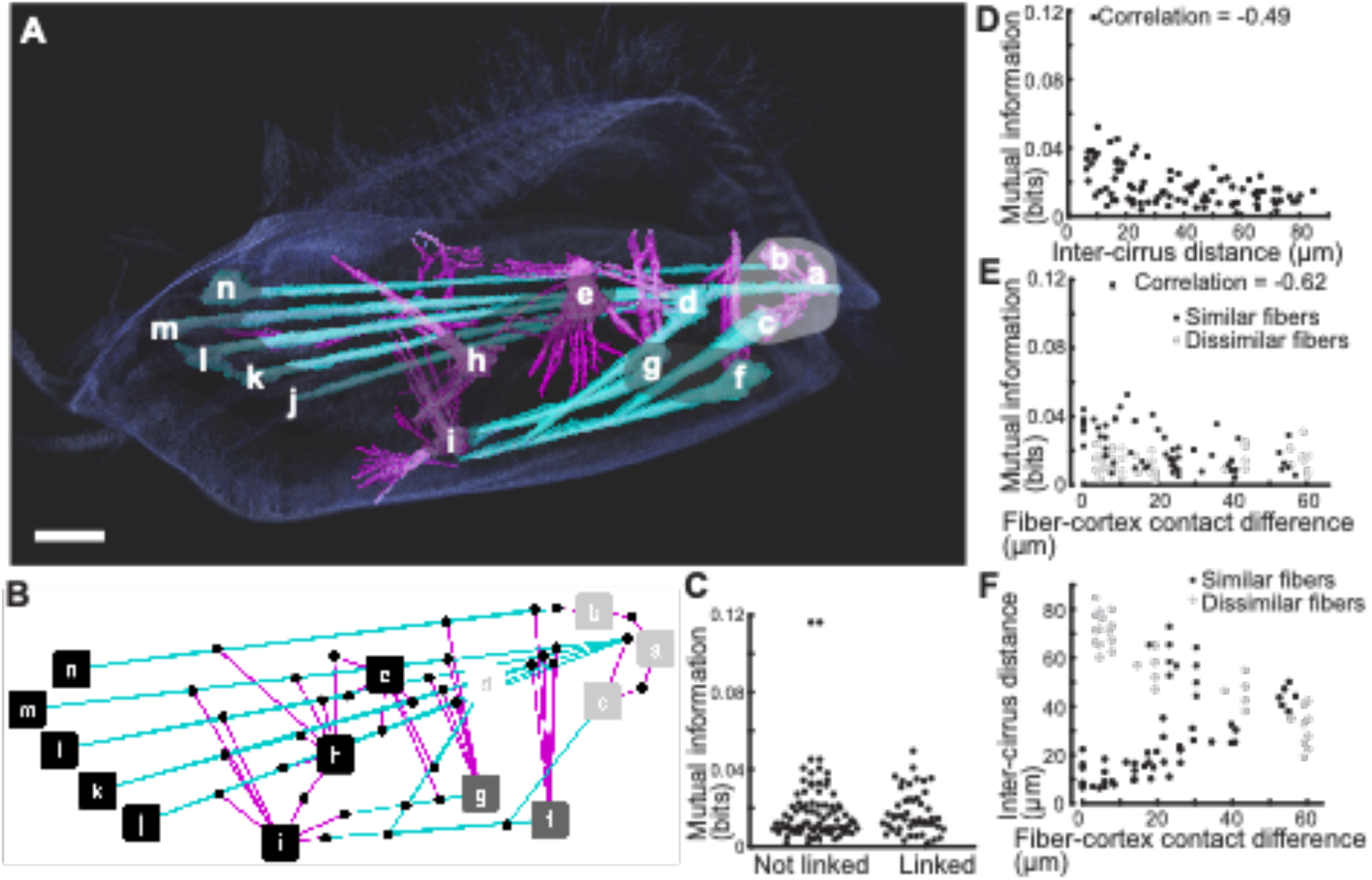
The structure of the complex, interconnected fiber system of *Euplotes* correlates with dynamical associations between cirri. (A) The SiR-tubulin labeled cell (faint, dark blue) was imaged by confocal microscopy, and a 3D reconstruction as obtained from serial confocal slices. Fibers were manually traced in each slice using TrakEM2 in FIJI. Two morphologically distinct classes of fibers were observed and are indicated as follows: thick, linear fibers are cyan and thinner, filamentous fibers are magenta (see Figure S3 for more image data). Fibers emanate from the base of each cirrus and form a connected network between all cirri. The base of each cirrus is indicated by corresponding letters (as in Figure 2C). Gray shading indicates the dynamical groups identified by dimensionality reduction and follows the same color scheme as in Figure 2H. Scale bar is 10 µm. (B) A graph representation of fiber-fiber connections illustrates the complex and interconnected nature of cirrus associated fiber topology. Nodes correspond to the cirri to which each fiber system is associated, and edges indicate connections between fiber systems. Colors of nodes indicate the same groups as in panel a, and colors of edges indicate the types of fibers connecting to one another, cyan for thick fiber connections, magenta for thin fiber connections, and purple for thick to thin fiber connections. (C) Pairs of cirri that are linked by fiber-fiber contacts show no statistically significant difference in mutual information compared to those lacking fiber-fiber contacts. The plot displays mutual information between all pairs of cirri grouped by the absence (Not linked) or presence (Linked) of associated fiber-fiber connections. Statistical significance was evaluated by the Wilcoxon rank sum test. Note that when pairs of cirri were grouped by fiber-fiber connection type, we did observe an increase in mutual information for cirri associated with only thin fiber-fiber connection and only thick fiber-fiber connections compared to those lacking fiber-fiber connections (see Figure S3). (D) A plot of mutual information as a function of inter-cirrus distance displays negative correlation, with a Spearman correlation coefficient of −0.49 (p<0.001). Plotted values are defined with respect to pairs of cirri. (E) A plot of mutual information as a function of fiber-cortex contact distance grouped by fiber type similarity and lack thereof displays negative correlation, with a Spearman correlation coefficient of −0.62 (p<0.001) for pairs of cirri with similar fiber types and no significant correlation for those with dissimilar fiber types. Similarity of fiber types is defined in terms of sharing at least some fiber types as defined in panel A. Fiber-cortex contact difference is measured by the mean cross nearest neighbor distance (see Method Details) for all fiber-cortex contact points associated with each cirrus. The negative correlation values from the data plotted in panels D and E indicate that cirri that are closer to one another and also cirri with fiber-cortex contacts in nearby regions of the cell tend to have higher mutual information, and indeed cirri that are both close to one another and with similar patterns of fiber-cortex contacts display the highest mutual information. (F) A plot of fiber-cortex contact difference versus inter-cirrus difference (as in panels D and E) illustrates that nearby cirri tend to have similar associated fiber-cortex contacts, highlighting that nearby cirri with similar fiber-cortex contacts share the most mutual information. See also Figure S3.

We reconstructed in 3D the microtubule-based fiber system of *Euplotes* associated with cirri and lying just beneath the cell cortex ^38, 41, 42^. Upon inspection of our confocal reconstructions of SiR-tubulin labeled cells (Figure 4A, Figure S3), we noted the presence of two morphologically distinct classes of fibers, one thicker and linear and the other thinner, splayed, and less linear, consistent with previous observations (Figure 1C, ^38, 41, 42^). Fibers emanate from the base of all cirri, appear to intersect one another, and also connect to the cortex of the cell at various points. Some cirri were found to be associated with only thick fibers, while others have both or only thin fibers. Based on apparent fiber intersections and convergences, we found the fiber system forms a continuous network between all cirri, with the fibers associated with the base of each cirrus intersecting the fiber system of at least one other cirrus (Figure 4A,B).

Contrary to the long-standing standing hypothesis from the literature ^84^, the functional modules (groups of co-varying cirri) identified in our dynamical analysis were not exclusively linked by dense fiber intersections (Figure 4A,B) ^38, 42, 84^. In fact, connections between cirri are not generally associated with any statistically significant difference in mutual information (defined in terms of the information that the activation state of one cirrus has concerning the other) compared to unlinked pairs of cirri (p=0.14 by Wilcoxon rank sum test, Figure 4C). However, information flow became apparent when fiber-fiber links were grouped by type (i.e. thick to thick fiber, thick to thin fiber, or thin to thin fiber). Under this grouping, we found that pairs of cirri associated with only thick to thick fiber and only thin to thin fiber links have increased mutual information compared to those without links (Figure S3). Interestingly, we found that cirri nearby one another and connected by fibers to similar regions of the cell cortex shared the most mutual information (Figure 2C,G, 4D,E), suggesting that if the fibers play a role in cirral coordination, coupling may also be mediated by mechanisms involving the cirrus and fiber-cortex interface. Cirri d, e, h, i, for example, share very little mutual information with any of the other cirri, and fibers emanating from the base of these cirri contact the cell cortex and other fibers at various unique points. On the other hand, cirri g and f, which share more mutual information than any other pair, are associated with both thick and thin fibers terminating at similar regions of the cell cortex. Indeed, distances between pairs of cirri and cross nearest-neighbor distance ^87^ between paired sets of cirrus-cortex contact points both show significant Spearman correlations (−0.49, p<0.001 and −0.62, p<0.001 respectively) to mutual information (Figure 4D,E). These correlations indicate that mutual information between pairs of cirri tends to increase with proximity and also tends to increase with similarity between fiber-cortex contact locations. Thus, the cirri with the highest mutual information are those that are close together with similar fiber-cortex connections (Figure 4D-F).

Together, these observations suggest a mechanism of mechanical coordination in which microtubule bundles allow groups of cirri to influence successive behavior of other groups of cirri. We first sought to test this hypothesized mechanism by perturbing the fiber system using drug treatments. We observed that nocodazole, a drug that blocks the polymerization of microtubules, affected walking motility, causing cells to walk along more compact trajectories due to increased turning (Figure 5A,B, Videos S3 and S4) in a manner reminiscent of historical reports of altered walking due to microsurgery in ciliate *Stylonychia* ^88^. In contrast to the effect of nocodazole, we found that the microtubule stabilizer paclitaxel caused cells to walk along less convoluted trajectories compared to controls (Figure 5C). Quantifying these effect from video microscopy of cells under darkfield illumination in terms of a scaled path length defined as the total integrated path length walked by cells scaled by the maximum radial distance traversed, where a decrease in linear runs decreases this scaled path length, we found that nocodazole significantly decreased and that paclitaxel significantly increased the scaled path length of cells compared to controls (Figure 5D). We further noted that nocodazole acted in a dose dependent and reversible manner (Figure 5D and Figure S4). To test whether the actin cytoskeleton might also regulate motility, we treated cells with actin inhibitors including latrunculin, cytochalasin, and jasplakinolide and observed no effect on motility (Figure S4). The specific and opposing effects of a microtubule polymerization inhibiting drug (nocodazole) versus a microtubule stabilizing drug (paclitaxel) were consistent with the microtubule cytoskeleton playing a key role in coordinating walking motility. Next, we checked whether nocodazole treatment had an observable effect on the fiber system. By analyzing 3D reconstructions of confocal images of SiR-tubulin labeled cells, we found that fiber length significantly decreased compared to controls in cells where microtubule polymerization was disrupted by nocodazole (Figure 5E-G). Further, we were unable to detect the presence of any thin fibers in four out of seven nocodazole treated cells used for fiber analysis. Of the three cells with detectable thin fibers, we never observed connections between fibers associated with cirri a, b, and c and any other fibers. Additionally, we did not observe any thin fibers making distal cortical contacts. We did note, however, that when thin fibers were visible, connections appeared to be the same as those in Figure 4A,B. We then investigated how cirral dynamics were affected. Following the gait annotation procedure detailed previously, we characterized the walking dynamics of 6 nocodazole treated cells for a total of 1133 timepoints. Of those timepoints, 681 corresponded to cirral configurations never observed in untreated cells with a total of 391 new unique configurations. Projecting these new configurations onto the NMF space we obtained previously, however, revealed that most of the cirral configurations in nocodazole treated cells were near or within the clusters corresponding to the gait states we obtained from untreated cells (Figure S4). This allowed us to map new cirral configurations onto the gait states (see Method Details). We did note that while close to the original gait states, new cirral configurations tended to skew towards more active cirri, and we also noted the presence of a new cluster involving movement in all or nearly all cirri, to which we assigned a new gait state (Figure S4). We also found that mutual information between cirri was higher in general, with many pairs of cirri sharing higher mutual information than the highest values obtain in untreated cells (Figure 5H, Figure S4). This increase in cirral activity and correlations is consistent with the fibers playing a role in conveying inhibitory information during unperturbed walking. In further support of this inhibitory role, we found that paclitaxel treatment also caused an overall increase in mutual information of pairs of cirri (Figure 5H).

**Figure 5.**
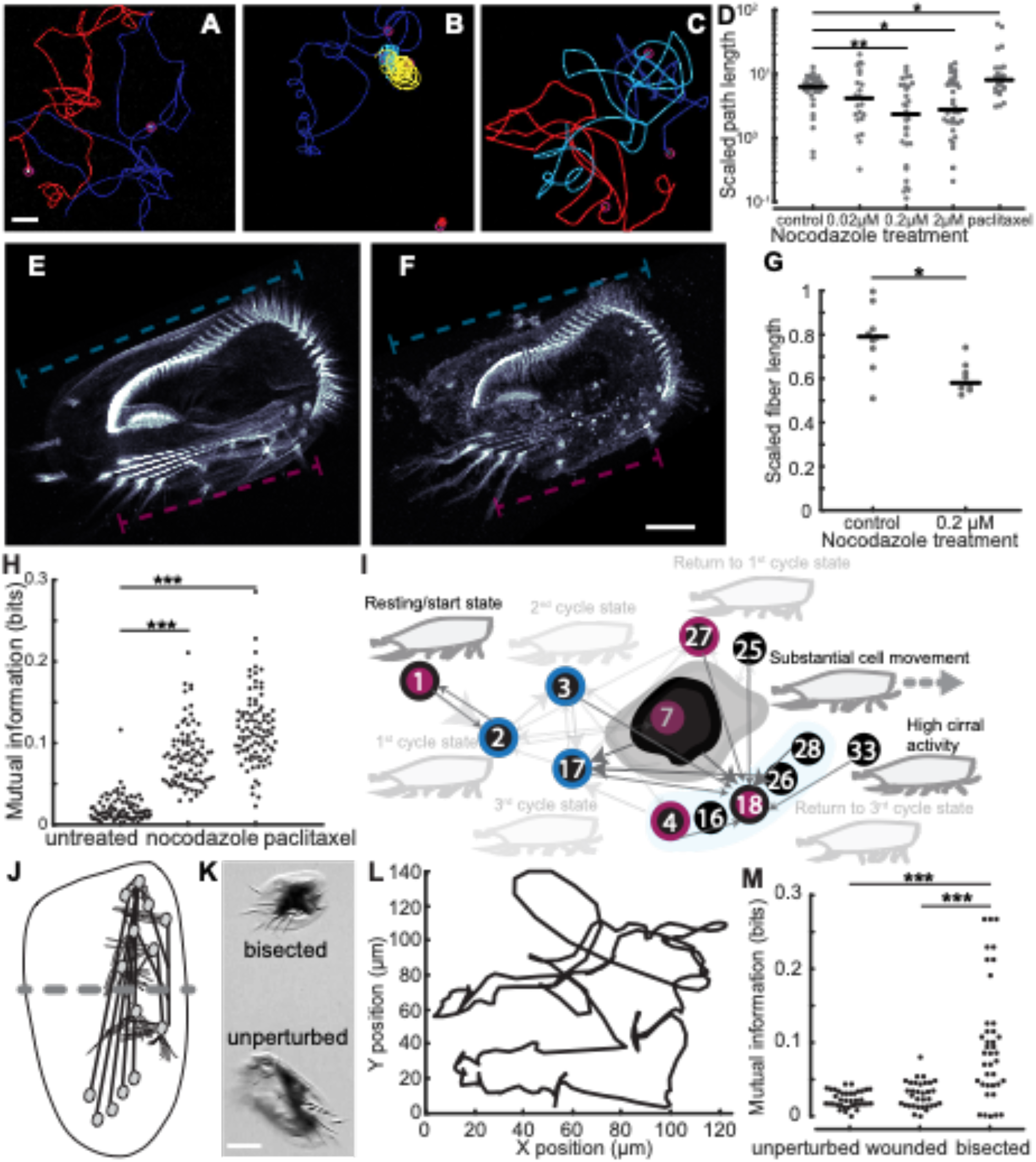
The fiber system of *Euplotes* mediates gait coordination. (A-C) Representative cell motility trajectories of cells imaged under darkfield illumination and tracked using the TrackMate plugin in FIJI ^108^ in control (A), nocodazole treated (B), and paclitaxel treated (C) cells highlight the increased turning causing curved and curled trajectories in cells treated with the microtubule polymerization inhibitor nocodazole and decrease in turns leading to smoother, less convoluted trajectories in cells treated with the microtubule stabilizer paclitaxel. Different colors represent the trajectories of different cells. Scale bar is 500 µm. (D) Nocodazole affects cell motility in a dose dependent and specific manner. Motility was quantified by the scaled path length, which is the total integrated distance walked by a cell scaled by the maximum distance the cell travelled from its starting point. This scaled path length decreases with a decrease in long, straight segments of trajectories corresponding to normal cell walking. Scaled path length decreased with increased nocodazole concentration, becoming significantly less than controls by 0.2 µM nocodazole. Note that at 2 µM nocodazole, cells were often observed to swim instead of walk, which may account for the lack of decrease in scaled path length compared to 0.2 µM treatment. In contrast to nocodazole, treatment of cells with 0.02 µM paclitaxel, which stabilizes microtubules, increases scaled path length compared to the control. The trajectories of at least 20 cells were analyzed for each condition. The black bars are median values. A single asterisk indicates p<0.05, and a double asterisk indicates p<0.005, as computed by a Wilcoxon rank-sum test. (E,F) Representative images illustrating the effect of the inhibition of microtubule polymerization by nocodazole on the fiber system and approach to quantifying this effect. Images are maximum intensity projections of confocal z-stack images of cells labeled by SiR-tubulin. Scale bar is 20 µm. (G) A plot showing that nocodazole treatment shortens fibers compared to controls. Scaled fiber length was measured by dividing the length of the longest fiber by the length of the entire cell. Nine cells were analyzed for each condition. The black bars are median values. A single asterisk indicates p<0.05 as computed by a Wilcoxon rank-sum test. (H) A plot of mutual information of all pairs of cirri shows that treatment of cells with the microtubule inhibitors nocodazole and paclitaxel change the distribution of mutual information due to an overall increase in cirral synchrony compared to untreated cells. The data for untreated cells is that from Figures 2 and 3. Nocodazole treatment was 0.2 µM, and 6 cells over 1133 timepoints were analyzed. Paclitaxel treatment was 0.02 µM, and 6 cells over 441 timepoints were analyzed. Three asterisks indicates p<0.001 as computed by a two sample Kolmogorov-Smirnov test. (I) A representation of the changes in the nature and organization of functional states as well as transitions between them highlights the effects of nocodazole treatment. This panel is partially adapted from Figure 3H and was produced by following the same analysis procedure. Any functional states and transitions depicted in Figure 3H no longer observed under nocodazole treatment appear faded. Arrows that are not faded represent the new highest probability transitions associated with the states. Outer colors of the nodes corresponding to the numbered gait states denote the type or absence of broken detailed balance associated with the gait state for untreated cells while the core color represents that for the treated cells. As in Figure 3H, black denotes the lack of unbalanced transitions, red denotes the state is only a source or only a recipient of unbalanced transitions, while blue denotes states which are both sources and recipients of unbalanced transitions. Note the loss of the cycle states as well as the functional states directing return to the cycle states, the emergence of the high cirral activity state involving activity in nearly all cirri, the reduction in gait states associated with the cloud of states of substantial cell movement (indicated by the reduction in the size of the black blob), and the emergence of many high probability transitions to gait state 18. (J) A cartoon illustrates the location of the cuts made for microsurgery experiments (dashed gray line). (K) A representative image shows a recovered bisected cell and a fully intact, unperturbed cell for reference. Scale bar is 50 µm. (L) A plot of the manually tracked trajectory of a walking, bisected cell illustrates the unsteady, curving manner of cell movement. (M) A plot of mutual information of all pairs of cirri h-n for unperturbed, wounded, and bisected cells demonstrating a change in the distribution of mutual information only in the case of bisected cells where fibers associated with h-n have been severed. Three asterisks indicates p<0.001 as computed by a two sample Kolmogorov-Smirnov test. The data for unperturbed cells is that from Figure 2, and data for wounded and bisected conditions involved 10 cells over 1308 timepoints in the case of the wounded cells and 1815 timepoints for bisected cells. See also Figure S4, Figure S5, and Videos S3-S7.

Next, we investigated how the dynamics of gait state transitions were affected. Following our previous analysis, we evaluated gait state transitions for the presence of broken detailed balance, separating state transitions into balanced and unbalanced transitions and found that the structure of state transitions differed greatly from that of unperturbed cells. Although some gait state such as 1, 2, and 17 were involved in high frequency transitions in both conditions, gait state transitions of cells with perturbed fiber systems exhibited less broken detailed balance and closer to equilibrium-like dynamics as indicated by an entropy production rate of 0.1 nats (compared to 0.4 nats in the unperturbed case), including the loss of the unbalanced, cyclic transitions (Figure S4). Figure 5I summarizes the change in the structure of gait state transitions including changes in broken detailed balance, reduction in transitions toward the states previously involved in cyclic transitions, and the reduction in the occupancy of states associated with the cloud of states involved in substantial cell movement, with only 10 of the original 21 cloud states sampled. We also noted that many of the new highest probability transitions feed gait state 18 (Figure 5I, Figure S4), which involves nearly exclusive activity in cirri f and g located close to one another at the edge of the cell. A persistent bias toward activation of these cirri, which are farthest from the central axis of the cell, may in part explain the increase in turning in trajectories. These results, in conjunction with the fact that cell velocities were indistinguishable control conditions (Figure S4) are consistent with predictions from simulations in which shuffling the order of gait state transitions reduced processivity without affecting speed (Figure 3N,O).

Although the result that disrupted gait coordination involves increased mutual information between cirri, note that this increase is due to more frequent joint activation or synchronous movements among cirri and so stems from reduced complexity of gait dynamics. For a complementary perspective linking these changes in gait dynamics to an underlying computational process, we applied the Causal State Splitting Reconstruction (CSSR) algorithm to construct *ϵ*-machines corresponding to walking cells ^48, 89^. *ϵ*-machines are automaton models consisting of a set of causal states with transitions between them and represent the minimal model consistent with accurate prediction of a stochastic process ^48^. The causal states of an *ϵ*- machine indicate how the process from which it is constructed stores information, and state transitions indicate how the process transforms information ^48^. We found that *ϵ*-machines constructed from the cirral activity of untreated cells tended to be similar to one another and were more complex in terms of having more causal states and transitions than those constructed from nocodazole treated cells, which also tended to be similar to one another (Figure S5, Method Details). This reduction in complexity may reflect reduced computational capacity of the nocodazole treated cells. Further, in conjunction with the decrease in entropy production under nocodazole treatment, these results are consistent with recent findings linking increases in broken detailed balance to increases in information processing in biological systems ^90^.

As a final additional test of the role of the fibers in mediating gait coordination, we revisited historical microdissection experiments. Although Taylor first reported in 1921 results of microdissection experiments indicating a role of the fiber system in mediating gait coordination ^84^, a reproduction of these experiments in 1966 with improved physiological conditions and quantitative analysis failed to observe disrupted coordination among cirri upon bisection of cells^85^. Importantly, neither of these reports involved analysis of walking behavior. We performed microdissections on cells using pulled quartz microneedles, severing cells transversely just in front of cirrus h, ensuring that we had severed all fibers associated with cirri j-n (Figure 5J,K). Similar to previous reports, we found that cell fragments regained spontaneous cirral activity after a brief recovery period. After 24 hr, we found that anterior portions of cells began to exhibit spontaneous walking activity (Figure 5K,L, Video S6), which persisted for up to 72 hr subsequently and has not been previously reported. Although these fragments were able to walk across surfaces, in all cases, they walked along curling or circular trajectories reminiscent of nocodazole treated cells except in the reverse direction (Figure 5L, Video S6). We analyzed the cirral dynamics of 10 fragments and found significantly higher mutual information shared between cirri indicating more synchronous cirral activity, similarly microtubule inhibitor treated cells, compared to cells that were merely wounded and displaying apparently normal motility (Video S7) or unperturbed (p<0.001 in both cases by Wilcoxon rank sum test), which displayed values consistent with one another (Figure 5M). As with the microtubule inhibitor experiments, these results are consistent the fibers playing an inhibitory role in gait coordination.

Taken together, these results are consistent with our hypothesis that the fiber system mechanically mediates gait coordination. Further, these results provide additional evidence that proper, active coordination of cirral dynamics and emergent gait states and gait state transitions is required for proper cellular walking.

## Discussion

Faced with the challenge of accounting for the emergence of apparently sophisticated cellular behavior, the directed walking of *Euplotes* driven by a seemingly disordered gait, we conceptualized the cell as a finite state machine. Traditionally, studies of computational processes performed by cells have focused on combinatorial logic, where the output of a computation depends only on the current input, performed by networks of molecules in cells ^25, 31–33^. We have focused on sequential logic, where outputs depend on the system state as well, an equally important aspect of the theory of computation with notable yet less developed representation in studies of cellular dynamics ^16, 20, 91^. Automata theory, which includes finite state machine models and necessarily involves sequential logic, provides tools for understanding structure and stereotypy in transitions between dynamical states, increasingly appreciated as features of the behavior of eukaryotic cells. Our approach revealed modularity in cellular dynamics associated with structural modularity of the cell (Figure 2, 4) in addition to stereotyped patterns of sequential activity (Figure 3) and yielded new insight into regulatory mechanisms (Figure 3, 5).

Related approaches involving the coarse-graining of complex dynamics and subsequent analysis have proved useful in revealing simplicity and stereotypy in the behavior and movement patterns of various organisms including flies, worms, and protists^3, 5, 65, 66, 92^. Reducing the complexity of biological dynamics in this way can be an important step toward making quantitative predictions and for interpreting the effects of perturbations. In general, particular approaches to coarse-graining will depend on details of the biological system, type of data collected, and underlying patterns in the data. The specific steps we have taken in the present study are best suited to biological processes that can be characterized in terms of discrete events. The analyses following coarse graining stand to provide the most insight into processes involving information processing where the sequential order of events may be important to the resultant function. Additionally, our perturbative experiments demonstrated how these approaches can drive mechanistic insight by helping connect function and computation to underlying mechanics and physiology.

Although there are examples of locomotor coordination reminiscent of the stochastic, non-equilibrium gait dynamics of *Euplotes*, such as gait switching in Kangaroo rats ^52^ or, most saliently, gait switching in an octoflagellate ^5^ and motility in cultured mammalian cells ^44^, walking locomotion in *Euplotes* represents a departure from many of the best studied appendage-based locomotor systems. For example, limbed locomotion in animals tends to proceed by highly stereotyped, determinate patterns of activity ^58, 62^, and many small, aquatic animals exhibit periodic movements of appendages, often cilia, during locomotion ^7, 63, 93^. Many forms of unicellular locomotion involve such dynamics as well including in sperm cells ^94^, diverse flagellates with various numbers of flagella ^63^, and ciliates ^63, 95, 96^. Even in cases where cellular locomotion involves fundamentally stochastic dynamics such as in run-and-tumble motility in *E. coli* ^12^ or analogous behaviors observed in protists ^11, 97–99^, motility can be described by equilibrium processes ^5^, in contrast to the non-equilibrium character of the gait of *Euplotes*.

Here, broken detailed balance in gait state transitions revealed cyclic activity with a mixture of stereotypy and variability in the gait of a single cell (Figure 3). To explain how these dynamics give rise to directed walking, we propose a mechanism, captured by our biophysical model, in which biased, actively controlled cyclic transitions serve to establish strain, effectively storing stress, in certain cirri, and the spontaneous release of these cirri from the substrate, during a series of unbiased gait state transitions, allows the cell to move forward. The cloud of unbiased transitions associated with substantial cellular movement is consistent with motility generation not depending on the precise order in which the strained cirri are released from the substrate. Return to the cycle states then are necessary to establish this process anew by winding up the system for continued, proper cell movement. Disruption in this resetting may lead to defects in walking as predicted by simulations (Figure 3K-O, 5) and with the reduction in entropy production and loss of the cycle states in irregularly walking cells with disrupted fiber systems, which appeared unable to consistently maintain linear trajectories (Figure 5B,D,I, Video S4). We find additional experimental support for our proposed mechanism in previously reported observations of cyclic velocity fluctuations in the trajectories of walking *Euplotes* ^36^.

We argue that subcellular processes are involved in actively coordinating cirri and propose that broken detailed balance in the gait of *Euplotes* indicates this active coordination. The results of experiments perturbing the tubulin-based cytoskeletal fiber system are consistent with its role in mechanically mediating communication both among cirri and between cirri and the cell cortex (Figure 4,5). We conjecture that movement of cirri relative to one another can establish tension in the fiber system and that the tension state of fibers associated with each cirrus may then modulate cirral activity in a manner reminiscent of basal coupling in flagellates^22^. Further, It is possible that microtubules may mechanically mediate the coordination of cellular processes in other eukaryotes, and microtubules in cells have recently been shown to respond directly to mechanical forces ^100^. Microtubules have also been shown and proposed to be involved in more complex signal transduction pathways^101, 102^. Changes in fiber length, as observed under nocodazole treatment, change the relative positions of cirri, which could lead to different distributions of tension and thereby induce the altered pattern of state transitions and associated walking defects. Our results show that perturbation of the fiber system shift the gait of *Euplotes* from a regime of asynchronous yet coordinated movement to a dysregulated regime with synchronous yet improperly coordinated movement. It should be noted, however, that the fact that cells with perturbed fiber systems, including bisected cells, display some walking ability implicates additional robustness and complexity in mechanisms of gait coordination and cell motility. Although details of the mechanism remain to be fully explored and tested, our results are consistent with the fibers system playing an active and inhibitory role in gait coordination. Thus, by combining information processing to properly dictate patterns of cirral activity and the mechanical actions of cirral movement, walking *Euplotes* embodies the sequential computation of a finite state machine. Furthermore, our approach to understanding a complex cellular behavior, grounded in theory of computation, allowed us to derive mechanistic insight from observational and perturbative experiments. Because reproducible biological function often emerges from the productive management of stochastic fluctuations, we expect our conceptual and analytical approaches may be useful in studying other living systems.

Our work lays a foundation for studies of sensorimotor behavior in *Euplotes*, which we believe will shed light on principles of cellular behavior. We have focused here on a single behavior (spontaneous, linear walking), but *Euplotes* exhibits a complex behavioral repertoire in response to diverse environmental cues, contexts, and physiological states^39, 40, 103104^. Many of these behaviors involve different patterns of cell motility. Our biophysical model of cell walking could help predict and interpret how changes in gait state transitions or the emergence of new gait states might give rise to these different patterns of movement. We speculate that the unusual gait of *Euplotes* and associated asymmetric cell structure, contrasting sharply with deterministic appendage movements and symmetric structure in animals and many swimming flagellates and ciliates, may reflect physical and phylogenetic constraints. The framework we have developed integrates across scales of space, time, and biological organization to link mechanics and subcellular dynamics and structure to overall cell movement patterns. Therefore, our work provides a way to begin to address the functional significance of cell morphology, gait patterns, and mechanisms of gait control, and in conjunction with comparative approaches, could provide new insight into how cellular behavior evolves.

Among the domains of life, eukaryotes uniquely display remarkable complexity and diversity in cellular behavior ^105^. Our approach, grounded in finite state machine analysis, has revealed modularity and stereotypy underlying complex cellular behavior and has provided insight into regulatory mechanisms across scales of biological organization. Our results suggest that integrating approaches from theoretical computer science, non-equilibrium statistical physics, and cell biology stands to shed light on the regulation of cellular behavior in eukaryotes more broadly. By revealing principles of cellular behavior, the line of research established here stands to advance our ability to predict, understand, and even one day engineer cellular behavior across diverse eukaryotic systems.

## STAR Methods

### Resource Availability

#### Lead contact

Further information and requests for resources, data, and code should be directed to and will be fulfilled by the lead contact, Wallace F. Marshall.

#### Materials availability

This study did not generate new unique reagents.

#### Data and code availability

All data reported in this paper will be shared by the lead contact upon request.

All original code has been deposited at GitHub and is publicly available as of the date of publication. DOIs are listed in the key resources table.

Any additional information required to reanalyze the data reported in this paper is available from the lead contact upon request.

### Experimental Model and Subject Details

#### Cell lines

Cultures of *Euplotes eurystomus* were obtained from Carolina Biological Supply Company (Item #131480) and were kept at room temperature under ambient light conditions.

### Method Details

#### Cell husbandry

Individual cells were isolated from cultures, which contained other protists and meiofauna, by pipetting and placed in non-treated 6-well plates (Thermo Fischer Scientific 08-772-49) containing spring water taken from cultures. Cells were kept in wells for no longer than five days before imaging, and if cells were to be kept for longer than 48 hours, wells containing cells were supplemented with 1% Cereal Grass Medium ^109^ (Thermo Fischer Scientific S25242) to prevent depletion of prey bacteria and otherwise maintain *Euplotes* under constant growth conditions.

#### Live cell brightfield microscopy

Cells were concentrated by centrifugation (500×g for 5 min) and resuspended either in 0.5 mL of spring water in coverglass bottomed FluoroDishes (World Precision Instruments FD35-100) or in 0.2 mL spring water on a coverslip (FisherScientific, 12-545-D) for imaging. No more than three cells were kept in 0.5 mL imaging samples and only one cell was ever kept in 0.2 mL imaging samples in order to minimize cell-cell interactions. Cells were observed to exhibit spontaneous walking activity on coverglass. Walking cells in FluoroDishes were imaged under brightfield illumination using a Zeiss Z.1 Observer and Hamamatsu Orca Flash 4.0 V2 CMOS camera (C11440-22CU) with a 20x, 0.8 NA Plan-Apochromat (Zeiss) objective. Cells on coverslips were imaged under brightfield illumination with coverslips inverted over a well containing a small amount of distilled water to reduce evaporation using a Zeiss Axio Zoom.V16 and a PCO pco.dimax S1 camera. Importantly, in both imaging systems, the focal plane was set at the interface between cirri of walking cells and the glass surface upon which they were walking. Images were acquired at 0.033 seconds per frame with a 0.005 second exposure in order to capture all cirral dynamics during walking with minimal blur.

#### Quantification of walking dynamics

Movies of walking cells were viewed using FIJI ^110^. Movement of cirri, or lack thereof was clearly visible in each movie frame (see Figure 2A and Video S1). The dynamical state of each cirrus in each movie frame was manually annotated. For each frame, each cirrus received a label of “1” if the cirrus was in motion and “0” if the cirrus was not moving and in contact with the coverslip. Motion of cirri was evident in terms of a change in cirrus shape or tip position often in addition to blur due to motion during image acquisition or position out of the focal plane (see Figure 2A and Video S1). While only slowly walking cells were recorded, sometimes cells nevertheless exhibit brief, spontaneous departures from slow walking during the course of movie acquisition. Any frame in which the movement of the cell and/or cirri were too fast to be resolved, such as during spontaneous escape responses ^40^ (Video S2), was excluded from analysis such that some videos were split into a number of separate continuous sequences. Thus, each movie frame associated with a particular time point in the walking trajectory, with the exception of those excluded from analysis as described, yielded a corresponding 14-element binary vector encoding the motility state of the cell in terms of the movement of cirri. Cell movement was tracked using the manual tracking feature of the TrackMate plugin in FIJI ^108^. The center of each cell was used as the reference feature for tracking. We analyzed the walking dynamics of 13 different cells.

#### Dimensionality reduction

Dimensionality reduction was performed by non-negative matrix factorization (NMF) implemented in MATLAB release 2019b (Mathworks, Natick). NMF was chosen as a dimensionality reduction technique to allow us to obtain a reduced, sparse, and interpretable representation of walking dynamics. Because NMF derives non-negative factors, the basis vectors in NMF space correspond straightforwardly to patterns of cirral activity. NMF involves factoring data, *A*, an *n* by *m* matrix, into non-negative factors *W,* an *n* by *k* matrix, and *H*, a *k* by *m* matrix where the product *W*\**H* approximates *A*. To determine the appropriate number of dimensions or rank, *k*, that are necessary to accurately represent the data without overfitting, we performed cross-validation by imputation with random holdouts ^111, 112^, also implemented in MATLAB. We randomly held out 15% of our walking dynamics data, performed NMF for a given *k*, and then used the NMF reconstruction *W*H*, to update the missing data entries. This process of updating is known as imputation, and we repeated the imputation process 50 times, by which point the imputed values had stabilized, to obtain a final NMF reconstruction. We then computed the root mean squared residual (RMSR) between the final NMF reconstruction, *W*H*, and our dataset, *A*. We performed this entire process 100 times for each value of *k*. As is generally the case for NMF, we observed a monotonic decrease in reconstruction error with increasing *k* without performing the imputation procedure ^113^ (Figure S1). In contrast to this trend, we observed an increase in RMSR of imputed values with increasing *k* indicating overfitting ^111^ (Figure S1). We chose *k*=3 because this value was the highest value before a notable increase in imputation error (Figure S1), which would indicate overfitting ^111, 112^. Thus, our choice of rank 3 selects the lowest rank approximation that captures structure of the dataset without overfitting that structure. Further, our choice facilitated the visual inspection of the structure of data in the reduced dimensional reconstruction.

Finally, we noted that for our chosen value of *k*, due to the stochastic nature of the NMF algorithm, which involves a random initialization step, we obtained slightly different solutions for different iterations ^111^. In order to choose the best reduced dimensional approximation, therefore, we performed NMF 500 times and chose the particular solution corresponding to the lowest RMSR compared to our dataset.

#### Clustering

Clustering on the dataset obtained using NMF was performed by density-based spatial clustering of applications with noise (DBSCAN) algorithm ^70^ implemented in MATLAB release 2019b (Mathworks, Natick). Structure in NMF space was clearly visible (Figure 2E-G), and DBSCAN using a Euclidean distance metric, was initially chosen as a clustering method because it yielded qualitatively good partitioning of the data. The DBSCAN algorithm involves stochastic search within neighborhoods of a given radius ε around datapoints, and points with a minimum number of neighbors, *n*, within their neighborhood are grouped as belonging to the same cluster, leaving two free parameters to determine. We set ε by first using the clusterDBSCAN.estimateEpsilon function in MATLAB (release 2020b, Phased Array System Toolbox), which yielded a value of 0.15. We next set about determining the minimum neighbor number, *n*. To do so, we computed the average Silhouette coefficient, a commonly used measure of clustering quality that indicates how well-separated clusters are ^114^, for various values of *n*. The results of this analysis are plotted in Figure S1. Higher Silhouette coefficients indicate better clustering, and we found that a value of *n*=8 maximized the mean Silhouette coefficient (Figure S1). We also noted, however, that for this value, many datapoints were found to be outliers, not belonging to any cluster due to having too few points within a distance of ε. Figure S1 displays percentage of datapoints found to be outliers as a function of *n*. In order to avoid categorizing more than 5% of datapoints as outliers, we chose to settle on *n*=4, which does not have a significantly different mean Silhouette coefficient compared to any of the others in the range *n*=2-7. This choice was further supported by the fact that major clusters involving more than 5 datapoints identified with *n*=8 were also identified with *n*=4.

Although this set of parameters gave qualitatively and quantitatively reasonable clustering results, we sought to further refine our clusters and to further reduce the outlier datapoints. We noted the obvious partitioning of the NMF dataset into three groups along the H2 axis (Figure 2E). We found the previously determined parameter values to yield good clustering for the top and middle partitions (H2≤1.1 and 0.2<H2<1.1), with no outliers. For the lower partition (H2≤0.2), however, we found that we were able to improve clustering by using ε=0.1182. With this updated value, we found no statistically significant change in Silhouette coefficient and reduced outliers to 0%. The clusters obtained by this process constituted the identification of the 32 gait states. We note here that the problem of determining the true or optimal number of clusters is an unresolved problem ^115^, and we note that we have followed standard methods to determine cluster number, and we found that our key results do not depend sensitively on the precise number of clusters identified (see following section and Figure S1 for more details).

#### State transition analysis

Following dimensionality reduction and clustering to identify gait states, we proceeded to characterize state transition dynamics. For each cell trajectory, we identified all unique gait state transitions for a total of 1423 unique pairwise transitions over the cumulative 2343 video frames for 77.14 s of recording. We computed empirical transition rates between states as the total number of observed transitions divided by the total time of observation. In order to determine which transitions were balanced and which were unbalanced, we followed Chang and Marshall ^45^, and performed binomial tests of statistical significance. Assuming a system at equilibrium, with all transitions obeying detailed balance, we expect to observe some deviation from exactly reciprocal transitions and can calculate the probability of observing a given set of ratios given underlying probabilities of forward and reverse transitions. The binomial probability of observing a set of transitions with known forward and reverse probabilities is given by

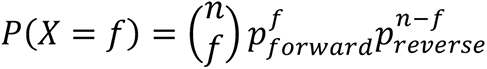

Where 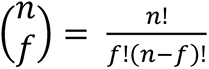 is the number of combinations, *f* is the number of forward transitions, *n* is the total number of transitions (such that *n-f* is the number of reverse transitions), and the probabilities *p*_*forward*_ and *p*_*reverse*_ are the forward and reverse probabilities. Considering only the set of transitions involving a specific pair of states, and calculating the probability that a transition between those states is either in the forward or reverse direction, the values of forward and reverse probabilities in the balanced case must be equal such that *p_forward_* = *p*_*reverse*_ = 0.5. With an α level of 0.05, we then considered reciprocal transition pairs with binomial probabilities less than 0.05 to be significantly unbalanced. Figure S1 displays the binomial probabilities associated with all transitions.

In order to calculate the estimated entropy production rate, we followed Wan and Goldstein ^5^, where the entropy production rate is defined as

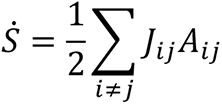

with conjugate fluxes *J_ij_* = *p_i_ k_ij_*− *p k_ij_* and forces 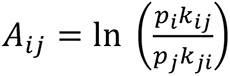 where the *p_l_* are the probabilities of being in state *l* at steady state and the *k_ij_* are the transition probabilities from states *i* to *j*. We estimate the state occupancy probabilities *p*_*l*_ as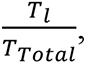, where *T*_*l*_ is the amount of time spent in state *l* over all trajectories and *T*_*Total*_ is the total recorded time, and the transition probabilities *k* as 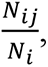, where *N_ij_* is the total number of observed transitions from state *i* to state *j* and *N*_i_ is the total number of transitions emanating from state *i*. To avoid *k_ji_* = 0 for pairs of states for which we did not observe any transitions during our experiments, we let 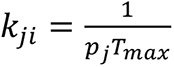 where *T*_*max*_ = 238 is the maximum number of observed transitions for any single recorded walking trajectory.

In the course of our state transition analysis, we also checked whether the waiting times between instances of each state might be non-exponentially distributed, with exponential distributions indicative of an embedded Markov process or possibly self-organized criticality ^116, 117^. Using the Lilliefors test implemented in MATLAB, we found that in general, waiting times were not exponentially distributed, although states 2, 3, 6, 16, 17, 18, 25, 27, 28, 32 were found to have waiting times consistent with exponential distributions with Benjamini-Hochburg corrected p-values of 0.046, 0.046, 0.022, 0.008, 0.046, 0.017, 0.046, 0.0081, 0.0046, 0.0046 respectively. Interestingly, none of the waiting times between the movements of individual cirri were found to be consistent with exponential distributions. These results are consistent with mechanisms constraining the temporal dynamics of cirri and state transitions.

In order to begin evaluating whether state transitions obeyed the Markov property for a discrete-time, first-order Markov process, where the transition probabilities from one state to the next are completely determined by current state ^75, 76^, we estimated the transition matrix for walking dynamics, consisting of the transition probabilities between all states. We estimate the transition probability from state *i* to state *j* as 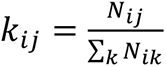 such that ∑*_i_ k_ij_* = 1. The entries of the transition matrix, *P*, are these transition probabilities with indices *i* for rows and *j* for columns. If gait state transitions obeyed the Markov property, we expect that the product of the transition matrix with itself, *P*^2^, would be equivalent to the two-step transition matrix where transition probabilities are computed as before except that state *j* is the state to which *i* has transitioned after an intervening transition. Figure S1 displays the results of this analysis showing that the two matrices show some quantitative and qualitative differences. Although these results strongly suggest violation of the Markov property, we applied the Billingsley test for a more statistically rigorous evaluation ^118, 119^. This test was implemented and performed in MATLAB. The Billingsley test gives a χ^2^ metric with *M*^2^*-*2*M* degrees of freedom given by

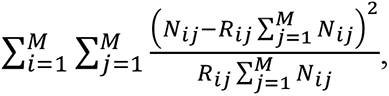

where *R*_ij_, the independent trials probability matrix, is given by

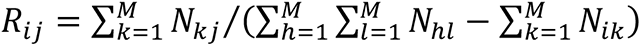. Applying this test to our gait state transition data, we found that the null hypothesis that the gait conforms to a first-order discrete time Markov process was rejected (p=0.005).

Importantly, we also noted that the key qualitative results of our state transition analysis are robust to the details of clustering results. In particular, we find that strongly unbalanced transitions and violation of the Markov property exist for a range of clustering parameters. Figure S1 displays the transition matrices for different clustering results.

To arrive at the simplified, state machine representation of the gait, we focused on the highest probability transitions emanating from each state. Transition probabilities were estimated as *k_ij_* (as defined above). This allowed us to prune away rare transitions in order to reveal the dominant structure of gait state transitions. Figure 3G displays the pruned transition matrix as a heatmap. We found that relatively few states were the recipients of the majority of high probability transitions, and many states received none. To more clearly visualize the structure of transitions, we grouped together all gait states receiving no more than one unique high probability transition based on the idea that state transitions into this group show little bias in terms of source state, and within the group, transitions between states exhibit low probability, time unbiased, equilibrium-like fluctuations.

#### Biophysical model and simulations

For our simple biophysical model, we consider a 2D system in which a *Euplotes* cell is walking across a surface in a low Reynolds number environment ^43^. The cell has 14 cirri, which exist in one of two states: actively moving or not actively moving, following our quantitative gait characterization. Cell position and orientation is defined in terms of equilibrium position of the cirri. Cirri can generate a motive force to drive cell motility when moving and resist displacement when not moving and in contact with the surface. For our model, we remain agnostic to the details by which cirri produce generate force noting only that in our experiments, no cell displacement was observed when cirri were not moving and in contact with the coverslip. We therefore let cirri generate a constant force in the direction dictated by cell orientation when moving. We conceptualize the resistance to displacement of unmoving cirri as stemming from the adhesive interaction between the cirrus and the substrate on which the cell is walking and the energy required to bend or deflect a cirrus. Consistent with experimental observations, we do not allow for translation of a cirrus-substrate contact point while a cirrus is not actively moving.

For a particle moving through a fluid at low Reynolds number, such as our cell, velocity 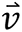 will be given by

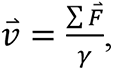

where 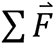 is the sum of the external forces acting on the particle, and *γ* is constant related to the geometry of the particle and the viscosity of the fluid ^120, 121^ accounting for drag. In our model, cirri are responsible for the forces involved in motility, so 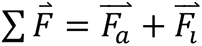 with the motive force 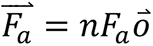 where n is the number of active cirri, *F*_*a*_ is the magnitude of the constant force generated by active cirri, and 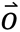 is the unit vector in the direction of cell orientation, and the resistive force 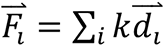 is a sum over the inactive cirri where k is a constant controlling the resistance of a cirrus to deformation and 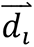 is the displacement vector of inactive cirrus *i*. We note that the forces driving cell motility in *Euplotes* stem from complex mechanical interactions, but for our model, we have chosen simple, first order expression to capture very basic features.

Similar to the expression for velocity above, angular velocity of a walking cell in our model is given by

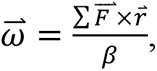

where *β* is a constant related to the geometry of the cell and viscosity of the fluid, 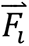 is the force due to cirrus *i* (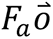 for active cirri and 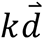 for inactive cirri), and 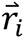 is the vector pointing from the center of the cell to cirrus *i*.

In addition to the relative positions of the cirri and patterns of cirral activity, the four parameters *γ*, *β*, *F*_*a*_ and *k* govern cell motility in our model. From these four parameters, we obtain three related dimensionless parameters:*α* = *β*⁄*γl* where *l* is the maximum distance between cirri, which can be thought of as characterizing the unsteadiness of the cell the intrinsic susceptibility of the cell to turning due to cell geometry; Μ = *F*_*a*_⁄*kl*, which can be thought of as characterizing degree to which cirral activity will tend to induce cell movement in opposition to inactive cirri; and ℱ = *F*_*a*_*t*⁄*γl* where t is the duration of a timestep in simulations, which can be thought of as the strength of the cirral motive force relative to the viscous drag experienced by the cell due to its fluid environment.

For all simulations, relative equilibrium cirral positions calculated from the average cirral positions in a video frame with no cirral activity over the 13 cells used for gait analysis (Figure 2) were used. In each simulation timestep, defined by the timestep used for recording videos for gait analysis, velocity and angular velocity are calculated based on the positions and activity of cirri, and the positions of all cirri are updated accordingly before proceeding. To calibrate the parameters of the model, we used the cirral patterns recorded from the cells used for gait analysis. We swept parameter space and found that simulations qualitatively and semi-quantitatively recapitulated experimentally measured cell motility with *α* = 0.001, Μ = 0.26, and ℱ = 0.008 (Figure 3K, Figure S1). These parameter values were subsequently used for all simulations.

For simulations with shuffled gait state transitions, we used MATLAB’s shuffle command on the cirral dynamics of actual cells to obtain sequences of gait state transitions of the same length as those that were experimentally obtained except in a random order. To obtain random patterns of cirral activity similar to those measured experimentally, we generated cirral activity according to a process defined by two probabilities: *p_a_*, the probability of transitioning from inactive to active at each timestep and *p_i_*, the probability of transitioning from inactive to active at each timestep. We initialized sequences with no cirral activity and then updated cirral activity according to these probabilities for each timestep in the sequence. We found that setting *p_a_*=0.3 and *p_i_*=0.1 yielded the same average cirral activity as that recorded experimentally, 0.23 ± 0.42 per frame and 0.24 ± 0.43 per frame respectively where the values are mean ± standard deviation. All simulations were performed in MATLAB.

#### Confocal microscopy

Cells were prepared for imaging and placed into a FluoroDish as described in the Live Cell Brightfield Microscopy section. Cells were then labeled with SiR-tubulin (Spirochrome provided by Cytoskeleton, Inc, CY-SC002) at 1 µM concentration. Cells were imaged using a Zeiss LSM 880 AxioExaminer and a 40x, 1.2 NA C-Apochromat water immersion objective (Zeiss) and excitation provided by a 633 nm laser (Zeiss). Only one full confocal z-stack of a complete cell was obtained during imaging to avoid effects of photodamage.

#### Fiber reconstruction and analysis

The image stack resulting from confocal imaging was first aligned in FIJI using the StackReg plugin ^110^. Next, fibers were manually segmented in each of the aligned z-stack images using the TrakEM2 plugin in FIJI ^122, 123^. Thick and thin fibers (Figure 4A) were morphologically distinguished, with thick fibers having a diameter of no less than 5 µm at the thinnest point. Fibers were traced from their distal termini to their convergences at the base of the cirri with which they were associated. Following segmentation, 3D surfaces were reconstructed in TrakEM2. Inter-fiber contacts were then found by inspection of 3D reconstructions and verified by examining individual z-stack frames to confirm intersections between fibers.

#### Drug treatment experiments and analysis

For all cytoskeleton inhibitor treatment experiments, 1 mL of cells in culture were placed in wells of 12-well plates (Thermo-Scientific, 12-565-321). Nocodazole (Sigma-Aldrich, M1404) as a stock solution of 6.64 mM in DMSO diluted further in distilled water, was added to achieve appropriate concentrations, with no more than 1 µL of additional volume added, and 1 µL of distilled water with equivalent DMSO concentration to nocodazole treatments added to controls. Cells were incubated for 1 hr before initiating experiments. No cell death was observed at any concentration of nocodazole in the 6 hours following nocodazole treatment. Paclitaxel (Sigma-Aldrich, T7191) as a stock solution of 2.23 mM in DMSO diluted further to 20 µM in distilled water, was added to achieve a final concentration of 20 nM to cells in solution. Latrunculin B (Thomas Scientific, C834E37) as a stock solution of 1.1 mM in ethanol was further diluted in distilled water and added to achieve a final concentration of 10 µM with cells in solution. Cytochalasin B (Fisher Scientific 1493-96-2) as a stock solution of 2.1 mM in ethanol was further diluted in distilled water was added to achieve a final concentration of 50 µM with cells in solution. Jasplakinolide (Fisher Scientific 42-012-750UG) as a stock solution of 1 mM in DMSO was further diluted in distilled water to achieve a final concentration of 10 µM with cells in solution. No cell death was observed in the 6 hours following treatment with any of the actin inhibitors. For the control condition for actin inhibitor experiments, both DMSO and ethanol was added to match the concentrations added in the cytochalasin B and jasplakinolide conditions.

##### Washout experiments

For nocodazole washout experiments, motility assays (described below) were also performed after placement of cells into well plates and before nocodazole treatment. Following 0.2 µM nocodazole treatment and another motility analysis, cells were picked in 5 µL of media and placed into wells of 6 well plates (Corning, CLS3736) each containing 2mL of fresh media. No more than 20 µL of nocodazole treatment media was added to any well so that the resultant nocodazole concentration in the washed condition was no more than 2 nM. Cells were allowed to recover in this condition for 4 hr, and then a final motility assay was performed.

##### Motility analysis

For motility analysis, cells were picked from well-plates and placed onto well-slides created by using a paper hole punch tool to punch a hole in 0.25 mm thick silicone spacer material (CultureWell, 664475), and adhering the spacer material to a glass slide (Corning, 2947-75X25). A total volume of 20 µL including up to 4 cells was added to well slides. A glass coverslip (FisherScientific, 12-545-D) was placed atop the well slide, sealing the well and creating an imaging chamber. After creating the imaging chamber, cells were allowed to acclimate to their new environment for 10 min. Cells were then imaged on a Zeiss Axio Zoom.V16 microscope under darkfield illumination with a Canon EOS T5i DSLR camera recording at 30 fps for 2 min. Videos were then processed using FIJI ^110^. First, images were cropped to remove extraneous parts of the field of view that did not contain the imaging chamber, and then background subtraction was performed by creating an image composed of mean pixel intensity values over all frames of the video and subtracting this mean image from all frames of the video. A mean filter with a four-pixel radius was then applied to each frame of the video for the purpose of smoothing. After processing, tracking was performed using the TrackMate FIJI plugin ^108^. For detection of objects (cells), a Laplacian of Gaussian filter was applied with an estimated blob diameter of 25 pixels and threshold of 0.1. A quality threshold was set manually when necessary to filter out any detected objects that were not cells. The Linear Assignment Tracker with a linking max distance of 15 pixels, gap-closing max distance of 150 pixels, and a gap-closing max frame gap of 200 frames was then used to generate linked tracks (trajectories) of detected cells. Trajectories were analyzed in MATLAB. Scaled path length for each tracked cell was calculated by summing the length of all segments of the track and dividing by the maximum distance the cell traveled from its starting point.

##### Fiber length analysis

Just prior to confocal imaging, cells were washed by picking up to five cells in 10 µL volume each and placing into 1 mL fresh media in a 12-well plate (Thermo-Scientific, 12-565-321). Cells were then prepared for imaging as described in the Confocal Imaging section of Method Details. Confocal z-stacks were then loaded into FIJI and aligned using the StackReg plugin ^110^. Cell lengths were determined by finding the maximal distance between two points on the front and rear ends of the cell. Because of variability in the detectable fibers in nocodazole treated cells, only fibers associated with the rear cirri (j-n), which were visible in all cells, were used for analysis. All of these rear fibers were measured, and the reported scaled fiber length was obtained by dividing the length of the longest fiber by the corresponding cell length. In all cases, the fiber associated with cirrus m was the longest fiber.

##### Analysis of cirral dynamics in nocodazole and paclitaxel treated cells

Cells were prepared for imaging and imaged as described in the Live Cell Brightfield Microscopy section of the Method Details with the exception that a Canon EOS T5i DSLR mounted on a Zeiss Axio Zoom.V16 microscope was used to record movements. Additionally, video was recorded at 0.066 seconds per frame to avoid blurring and then videos were downsampled to 0.033 seconds per frame for analysis. Cirral dynamics were quantified as described in the Quantification of Walking Dynamics section of the Method Details.

To assign cirral configurations of nocodazole treated cells to previously identified gait states, we first matched any cirral configurations with known gait state identity. Next, due to proximity of new cirral configurations not observed in untreated cells (Figure S1), we were able to map the new cirral configurations onto the clusters defining the gait states by determining the nearest cluster to the new cirral configuration. Distance between new cirral configurations and clusters were determined by finding the shortest distances to points defining convex hulls of each cluster. The shortest of all these distances then indicated the nearest cluster and corresponding gait state to which the new cirral configuration was assigned. We noted an obvious dense cluster of points corresponding to activation of nearly all cirri, and we identified a new cluster and corresponding gait state by applying the DBSCAN algorithm as described in the Clustering section of the Method Details. Evaluation of transition dynamics was performed as described in the State Transition Analysis section of the Method Details. This analysis was all performed in MATLAB.

#### *ϵ*-machine construction

Our representation of the Euplotes at timestep t takes the form of a length 14 binary string 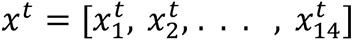 where 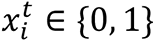. For reducing dimensions, we found that bigrams of the 14-dimensional strings, yielded more consistent, interpretable results than unigrams, so 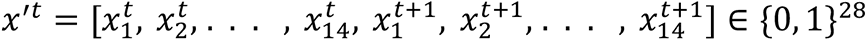. In order to learn latent states, we used Variational AutoEncoders (VAEs) ^124^ to reduce each fourteen dimensional timestep to three dimensions. VAEs in particular are used for their ability to learn an interpolatable latent space where high-dimensional training data points are mapped to points in low-dimensional space that mimic a normal distribution centered around the origin. This processes yields a smooth latent space with dimensions that represent core aspects of the data. We used a very minimal VAE with an Adam optimizer ^125^, consisting of one dense layer for the generating encoder means and one dense layer for generating encoder variance. This creates a 3-dimensional normal-like distribution, which we sample from and decode with a one layer decoder.

VariationalEncoder(*x*’^*t*^) → *z*^*t*^ ∈ ℝ^3^

After reducing the dimensionality of our data, we perform spectral clustering ^126^ using scikit-learn ^127^ with 100 k-means runs and 4 clusters, where the cluster count was decided by inspection of the resulting classes.

SpectralClustering(*z*^*t*^) → *c*^*t*^ ∈ {1, 2, 3, 4}

Applying this procedure to the data from the cells used for gait analysis (the same as used for Figure 2) with bigram training parameters lr=0.0011, weight_decay=0.000727, k=3, hidden_dim=5, beta1=0.5, and beta2=0.900, we found the following four clusters corresponding to particular patterns of cirral ativity: 1) Front cirral activity 2) High overall cirral activity 3) Rear cirral activity and 4) Little cirral activity (Figure S5). When we applied this procedure to the data from cells treated with 0.2 µM nocodazole (the same data used for Figure 5) with bigram training parameters lr=0.0026, weight_decay=0.000106, k=3, hidden_dim=3, beta1=0.5, and beta2=0.999, we obtained only three distinct clusters with similar cirral activity patterns to those of untreated cells except that the rear cirral activity cluster (3 above) disappeared (Figure S5). These clusters represent an independent, alternative coarse-graining of patterns of cirral activity into gait states to that presented in Figure 2. Although independent, this procedure identified the same underlying structure in the data. In particular, the alternative gait states here involve similar patterns of cirral activity to those defining the NMF components depicted in Figure 2H. Whereas the spatially distinct groups of cirri sharing the most mutual information define the space of gait states under the NMF-based approach, under this alternative, activity or complete lack thereof in those same groups of cirri defines the gait states. Furthermore, the loss of cluster 4 under nocodazole treatment is consistent with the loss of gait states as depicted in Figure 5I.

To construct *ϵ*-machines ^79^, characterized by a set of causal-states and transitions between them, we feed a stream of behavioral symbols *c*_P_, *c*_8_,. . . , *c*_*T*_ into a Causal-State Splitting Reconstruction (CSSR) ^89, 128^ algorithm. Due to the nature of the CSSR algorithm, we can only construct *ϵ*-machines from unbroken data streams, so we constructed *ϵ*-machines for each cell from a given dataset. Performing this process over gait state data streams derived from cirral activity recordings as described above for each cell from the untreated and nocodazole datasets yielded a set of *ϵ* machines. Representative examples of *ϵ*-machines obtained from each dataset are depicted in Figure S5.

##### Microsurgery experiments

All microsurgeries were performed by hand under observation with a Zeiss Axio Zoom.V16 using quartz microneedles pulled from quarts rods (Sutter Instrument, QR100-10) using a Sutter Instrument P-2000 Laser-based micropipette puller. Individual cells were picked and placed in a 5 µL droplet at the center of an imaging chamber as described in the Motility Analysis section of the Method Details. For bisections, microneedles were quickly and firmly pressed onto cells just anterior to cirrus h, cleanly severing the cell including the fibers associated with cirri j-n and sealing the cell membrane. Cells were allowed to recover for at least 5 minutes until some motility resumed, and then 15 µL of fresh culture media was added to the imaging chamber, which was subsequently sealed with a glass coverslip (FisherScientific, 12-545-D) to minimize evaporation. After 24 hr, once cell fragments resumed walking motility, gait analysis was performed as described in the quantification of walking dynamics section of the Method Details, except that cell fragments only had 7 cirri (j-n) instead of the full 14. Experiments with wounded cells were conducted in the same fashion as described for bisections except that instead of cutting cells in half, wounding was performed by stabbing with a microneedle a portion of the cell that does not have any fibers and was confirmed visually by the loss of some amount of cytoplasm.

### Quantifications and statistical analysis

Statistical details of the experiments can be found in the figure legends, the main text, or the Method Details section. Statistical details include exact value of n, what n represents (generally the number of cells measured), definitions of center, and dispersion and precision measures. Statistical tests and fits were performed as described in the Method Details section using MATLAB release 2019b or 2020b (Mathworks, Natick).

### Supplemental Video Captions

**Video S1**. **A walking cell.** A representative example of a single *Euplotes eurystomus* cell walking across a coverslip using motile cirri illustrates the walking gait. Note that the movement of cirri, or lack thereof is clearly visible in each frame of the movie. This cell was imaged using a Zeiss Axio Zoom.V16 microscope under brightfield illumination. The video is slowed down by a factor of 4 from real time to show that the movements of cirri are clearly visible.

**Video S2**. **Walking motility including escape responses.** A representative example of a single *Euplotes eurystomus* cell walking across a coverslip using motile cirri including escape responses, which were excluded from gait analysis. Escape responses occur at t= 18s, 20s, and 25s. This cell was imaged under brightfield illumination using a Zeiss Z.1 Observer with a 20x, 0.8 NA Plan-Apochromat (Zeiss) objective. The video is slowed down by a factor of 4 from real time so that the movements are clearly perceivable.

**Video S3. Cell tracking for motility analysis.** A representative example of the trajectories of cells imaged at low magnification under darkfield illumination using a Zeiss Axio Zoom.V16 microscope. Cells appear as bright objects with magenta circles around them. Different colored lines correspond to the tracks of different cells tracked using the FIJI plugin TrackMate.

**Video S4. Motility of nocodazole treated cells.** A representative example of the trajectories of cells treated with 200 nM nocodazole imaged at low magnification under darkfield illumination. Cells appear as bright objects with magenta circles around them. Different colored lines correspond to the tracks of different cells tracked using the FIJI plugin TrackMate. Note the confined movements of cells due to a decrease in long, linear runs. Compare to Movie 3 for the control condition.

**Video S5. Motility of paclitaxel cells.** A representative example of the trajectories of cells treated with 20 nM paclitaxel imaged at low magnification under darkfield illumination. Cells appear as bright objects with magenta circles around them. Different colored lines correspond to the tracks of different cells tracked using the FIJI plugin TrackMate. Note the increase in long, linear runs and decrease in abrupt changes in direction and tight turns. Compare to Movie 3 for the control condition.

**Video S6. A walking, bisected cell.** A representative example of a bisected cell analyzed in microsurgery experiments displaying the characteristic backwards, spiral walking motility defect. This cell fragment was imaged using a Zeiss Axio Zoom.V16 microscope under brightfield illumination. The video is slowed down by a factor of 4 from real time to clearly show the movements of cirri.

**Video S7. A walking, wounded cell.** A representative example of a wounded cell analyzed in microsurgery experiments. This cell was imaged using a Zeiss Axio Zoom.V16 microscope under brightfield illumination. The video is slowed down by a factor of 4 from real time to clearly show the movements of cirri.

## Acknowledgements

We would like to thank Nicole King for the use of the confocal microscope and Adair Oesterle for assistance with microsurgery. We would like to thank the students and faculty of the 2016 Marine Biological Laboratory Physiology Course and current and former members of the Marshall Lab and King Lab for comments, critiques, and encouragement during the development of this project. We would also like to thank David Booth, Greyson Lewis, Dennis Bray, and members of the Fourmentin-Guilbert Scientific Foundation for comments on the manuscript. This work was funded by the I2CELL Seed Award of the Fourmentin-Guilbert Scientific Foundation (WFM, JBP). Additional funding was provided by NIH grant R35 GM130327 (WFM), NSF grant MCB-2012647 (WFM), and a Merck Fellowship of the Jane Coffin Childs Memorial Fund for Medical Research (BTL).

## Author Contributions

BTL, JDP, and WFM conceived of the study. BTL, JG, and WFM developed methodology. BTL performed the experiments, and all authors analyzed and interpreted data. BTL and WFM originally drafted the manuscript, and all authors reviewed and edited the work.

